# Linking *in situ* activity to fitness reveals that clinically-relevant factors underpin pathogen survival in environmental reservoirs

**DOI:** 10.64898/2026.02.12.705432

**Authors:** Charles Nunez, Thomas D. Watts, Nguyen Thi Khanh Nhu, Jessica Solari, Thanavit Jirapanjawat, To Nguyen Thi Nguyen, Thuy Nguyen, Gareth Howells, Yue Qu, Luis E. Valentin-Alvarado, Steven J. Hancock, Yang Liao, Wei Shi, Trevor Lithgow, Mark A. Schembri, Chris Greening, Francesca L. Short

## Abstract

Natural environments are a key reservoir for many opportunistic pathogens, yet mechanisms enabling survival within complex microbial communities remain poorly defined. Current approaches do not adequately link molecular understanding of bacterial survival to the activities of bacteria within natural microbial communities. Here we combine *in situ* community profiling with isolate-level functional genomics studies to understand how opportunistic pathogens from the genera *Escherichia*, *Klebsiella* and *Enterococcus* survive in urban-adjacent freshwater ecosystems. Potential pathogens from these genera were identified and isolated from urban freshwater, and targeted metatranscriptomics showed expression of genes involved in housekeeping, resistance and virulence functions *in situ*. We further investigated an *E. coli* isolate of ST73, a clinically-important lineage associated with urinary tract infections, showing that it is pathogenic in mice, and survives in freshwater microcosms for at least one month. Functional genomics investigations in freshwater containing autochthonous microbiota revealed genes involved in amino acid metabolism, nucleotide biosynthesis, and biogenesis of the virulence- and biofilm-associated surface amyloid curli as important survival determinants; many of these genes were also highly expressed *in situ*. Deletion of curli structural genes resulted in reduced pelagic biofilm formation and a fitness defect specifically in the presence of freshwater microbiota, which was reversed by carbon supplementation, indicating that curli promote *E. coli* freshwater through aggregation-mediated nutritional competition. This study deepens our understanding of pathogen survival in waterways, highlights the importance of natural environments for selection of clinically-important traits, and provides a broadly applicable framework for interrogating mechanisms of pathogen survival in complex environments and communities.

**Significance Statement:** Many bacterial pathogens are not human-adapted, but are generalists able to thrive in environmental, animal- and human-associated niches. Understanding environmental survival could inform strategies to limit survival and break the transmission cycle. Here we pioneer a new approach to understand survival of pathogens within complex environments. Combining targeted isolations, in situ investigations and microcosm experiments, we show that antimicrobial-resistant pathogens are abundant and active within an urban water ecosystem, and identify the survival mechanisms of an E. coli strain from this environment. We reveal that curli, a virulence-associated surface structure, promotes environmental survival by enabling E. coli to compete with the native microbiota for nutrients. Our findings show that natural environments are selective niches shaping virulence traits in bacterial pathogens.

## Introduction

Many prevalent human pathogens are not truly human-adapted but are widely distributed microbes able to survive in diverse environments. Notably, Enterobacteriaceae family pathogens such as *Escherichia / Shigella* spp. and *Salmonella*, which collectively cause over a quarter of antimicrobial resistance (AMR)-associated deaths (1) and diarrheal disease that kills 500,000 children each year (2, 3), are frequently recovered from non-human niches such as soil and waterways (4). Natural environments, in particular waterways, are central to both the transmission and evolution of such pathogens because they provide a means for the bacteria to reach and colonise new hosts, equip the bacteria with beneficial genes through horizontal gene transfer, and select for clinically-relevant traits (5–8). Environmental monitoring has revealed a high prevalence of AMR Enterobacteriaceae in natural environments, in which *E. coli* is frequently used as an indicator of water quality (4, 9, 10). *E*. *coli* is an exemplar of opportunistic pathogens for which environmental survival is critical to success. *E. coli* exhibits enormous diversity encompassing commensal strains as well as pathotypes that cause intestinal and extra-intestinal infections (11, 12). *E. coli* can persist and disseminate in aquatic and terrestrial environments (13), and several pathogenic lineages including shiga toxin-producing enterohemorrhagic *E. coli* (e.g. O157:H7) and uropathogenic *E. coli* (UPEC) exhibit high rates of environmental persistence (14–17).

Despite this, we lack a systematic understanding of how opportunistic pathogens adapt to environmental reservoirs. Key studies in this area have revealed cell wall integrity is critical for *E. coli* K-12 survival in lake water (18), energy storage compounds enhance survival of *Vibrio cholerae* in pond water (19), energy metabolism supports survival of *Legionella pneumophila* in laboratory tap water incubations (20), among other factors. However, previous investigations have often relied on well-characterised laboratory strains rather than clinical or environmental isolates, targeted individual genes rather than taking a genome-wide approach, and/or employed laboratory rather than environmental conditions. One reason for these gaps is a limitation in the application of genome-scale methods to evaluate the determinants of pathogen adaptation to the environment. Yet modern genome-resolved meta-omic and functional genomics tools enable us to systematically understand what genes are expressed *in situ* and required in different environments (21–23). While these approaches are widely used to examine pathogen colonisation in hosts, they have yet to be combined to reveal how pathogens adapt to environmental settings. In the case of *E. coli*, survival in waterways and sediments has primarily been explored from the perspective of its utility as an indicator of faecal contamination, and has been shown to be affected by carbon and nitrogen availability, pH, salinity and temperature (17, 24–26). Though many of the genes and pathways involved in responding to these stresses are known through the collective molecular microbiology literature (27–35), few have been shown to contribute to survival *in situ*.

In this study, we aimed to gain systematic and mechanistic understanding of how pathogens adapt to their environment. To achieve this, we combined three key methodological innovations: (i) mapping isolate genomes to metagenomes and metatranscriptomes to understand gene expression of opportunistic pathogens in their environmental reservoirs; (ii) application of transposon-directed insertion-site sequencing (TraDIS) directly to environmental isolates to define genes required for their survival; and (iii) environmental microcosms to simulate how pathogens respond to different conditions and presence of resident microbiota. We focus on an urban freshwater environment in suburban Melbourne, Australia, from which multiple pathogens were isolated and shown to be transcriptionally active *in situ*. Functional genomic analysis of a mouse-virulent *E. coli* isolate (Ec19), from the ST73 lineage, revealed key pathways involved in survival and showed these are highly expressed in the environment, with the presence of the native microflora dramatically affecting fitness requirements. Our findings reveal new determinants of pathogen survival in freshwater and provide a framework to understand pathogen behaviour in complex environments.

## Results

### Antimicrobial-resistant opportunistic pathogens are present and active in urban-adjacent freshwater ecosystems

To understand how pathogens survive in urban-adjacent freshwater microbial communities, we combined genome-resolved metagenomic and metatranscriptomic analyses with isolation of putative opportunistic pathogens using four independent surface water samples (DC0 for the majority of pathogen isolation, DC1, DC2, DC3 for metagenomics and metatranscriptomics) from Dandenong Creek in Melbourne, Australia (**Figure S1A-C**). The most abundant and transcriptionally active taxa were Proteobacteria, Bacteroidota, Cyanobacteria, and various eukaryotes (e.g. SAR and Archaeaplastida), similar to previously described freshwater microbial communities (**Figure 1A; Figure S1D-F**) (36, 37). Putative opportunistic pathogens were also present and active, primarily within the families Enterobacteriaceae and Enterococcaceae which respectively comprised an average of 0.78% and 0.031% of the community (**Figure S1G**), with the genus *Escherichia* showing the highest relative transcriptional activity (**Figure 1A**). Metabolic profiling suggested most community members were obligate aerobes or facultative anaerobes with a broad capacity for organic carbon acquisition, light harvesting, and lithotrophy (**Figure S2A-B**). Antimicrobial resistance genes (ARGs) conferring resistance to aminoglycosides, disinfection agents (predominantly *qacG* genes) and rifamycins, as well as multidrug efflux pumps, were abundant and expressed (**Figure 1B, Table S1**). Bacterial virulence-associated factors from seven different categories (38) were also detected in each metagenomic and metatranscriptomic dataset: most highly expressed were secretion systems (including T2SS, T3SS T6SS and T4SS) and those for motility such as flagella (*Aeromonas spp*., *Pseudomonas aeruginosa, Burkholderia pseudomallei*) and iron-acquisition such as siderophores (Enterobactin, Aerobactin, followed by colonisation-related factors such as Type IV pili (*Aeromonas spp*., *P. aeruginosa*.) and Type I fimbriae (*Aeromonas spp.*) (**Figure 1C, Table S2**). These data show that members of this freshwater community not only have capacity for virulence and AMR, but actively express genes associated with these processes *in situ*.

**Figure 1.**
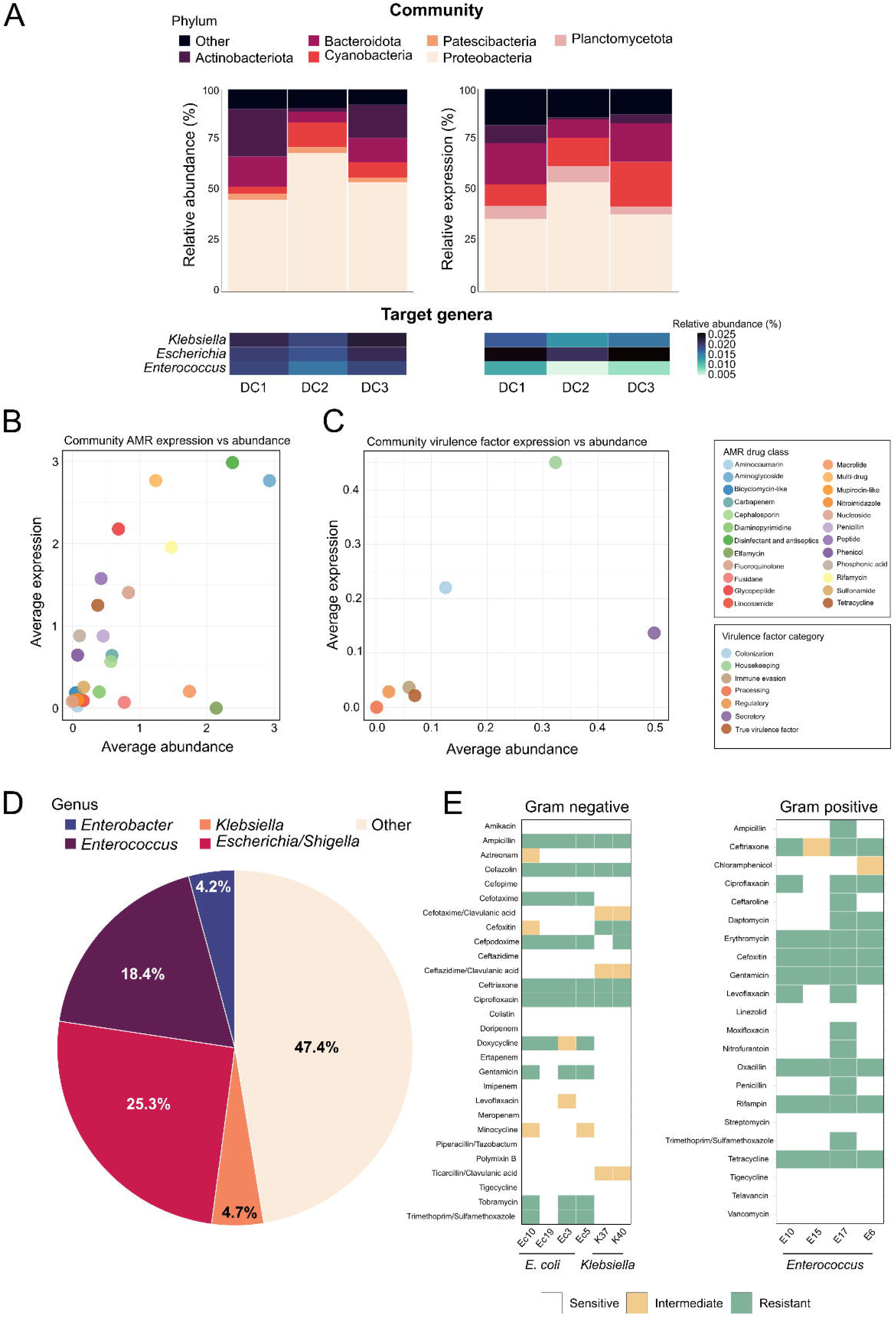
Freshwater microbial community composition, AMR profiles, and *in situ* gene expression. A) Stacked bar charts showing relative abundance and expression of top bacterial phyla in urban freshwater microbial communities, heatmaps below indicate average abundance and expression of target genera determined by mapping to isolate genomes. B) Expression vs abundance scatter plot of whole community AMR grouped by drug class. C) Expression vs abundance scatter plot of whole community virulence factors grouped by MetaVF categories. D) Results of targeted isolations from Dandenong Creek in Melbourne Victoria (see Supplementary Fig. S1) using chromogenic agar, with the percentage of each key genus indicated. E) Antibiotic susceptibility testing of Enterococcaceae (E10, E15, E17, E6) and Enterobacteriaceae (Ec10, Ec19, Ec3, Ec5, K37, K40) with ESBL, Gram-positive and Gram-negative Sensititre kits.

To isolate potential opportunistic pathogens, bacteria were cultured from water samples using chromogenic media, and provisionally identified by 16S rRNA gene sequencing. Recovery conditions were designed to be reasonably permissive (see Materials and Methods), but did not include specific measures to resuscitate viable but non culturable cells in the water samples, though the presence and activities of such cells would contribute to the community-level metagenome and metatranscriptome profiles. We identified 193 isolates from putative pathogenic genera (**Table S3, Figure 1D**), including 35 *Enterococcus*, nine *Klebsiella*, and 46 *Escherichia/Shigella*, with ten further characterised through whole genome sequencing and genotyping with pubMLST and Kleborate (39, 40) (**Table S4**). Several isolates were from clinically relevant lineages, including *E. faecium* sequence type 80 (ST80) that belongs to the globally disseminated *E. faecium* clonal complex 17 (CC17) (isolate E17), *E. faecalis* ST16 associated with nosocomial infections (isolates E6 and E10) (41, 42), and the emergent opportunistic pathogen *Klebsiella aerogenes* (isolates K37 and K40) (43, 44). An *E. coli* isolate was identified as ST73 (isolate Ec19), a clinically relevant lineage of extraintestinal pathogenic *E. coli* that commonly causes urinary tract infections (UTI) (45, 46). Isolates showed a range of resistance phenotypes (**Figure 1E**), with several resistant to multiple antimicrobial classes. The Enterobacteriaceae isolates exhibited extended-spectrum beta-lactamase (ESBL) activity and one *E. faecium* isolate was resistant to 16 antibiotics, including daptomycin.

To infer if these pathogens, or highly similar isolates, were active *in situ* and which genes they use to survive, we mapped whole community metatranscriptomic data to the individual isolate genomes within a hybrid index also comprising either the Enterobacteriaceae or Enterococcaceae metagenome contigs, and categorized them by COG categories and KEGG modules (**Figure 2A-D, Supplemental Note**). Enterobacteriaceae isolates showed the highest number of active processes, with genes from almost all COG categories active and represented. Enterobacteriaceae isolates were also more abundant and transcriptionally active than *Enterococcus* isolates based on RNA / DNA ratios, with Ec19 showing the highest relative transcriptional activity (**Figure 2A, Table S6**). Translation-associated processes dominated in all isolates, even following ribosomal RNA depletion (**Figure S3A, Table S5**), with high transcription of ribosomal RNAs, tRNAs, RNase P, SsrA, and core elongation factors. This was followed by expression of genes encoding RNA polymerase subunits. Aside from these core processes, genes involved in energy production and central carbon metabolism were expressed, including those encoding glyceraldehyde-3-phosphate dehydrogenase and acetyl-CoA carboxylase, genes encoding glycolysis or gluconeogenesis pathway enzymes, and ATP synthase components. Protein quality control and stress response associated genes *groEL/hsp60*, *clpAXP* and several cold shock proteins were also represented. Transcripts associated with the broad categories of cell division, envelope biogenesis, and curli production / biofilm formation were also detected, with genes encoding subunits of the biofilm-associated amyloid curli showing higher expression (**Figure 2C**). The *E. coli* isolates expressed genes for synthesising the siderophore aerobactin (*iucC*) and enacting catabolism of various amino acids, sugars, and other metabolites (e.g. D-galactonate), suggesting that they can scavenge multiple resources in nutrient-limited freshwater environments.

**Figure 2.**
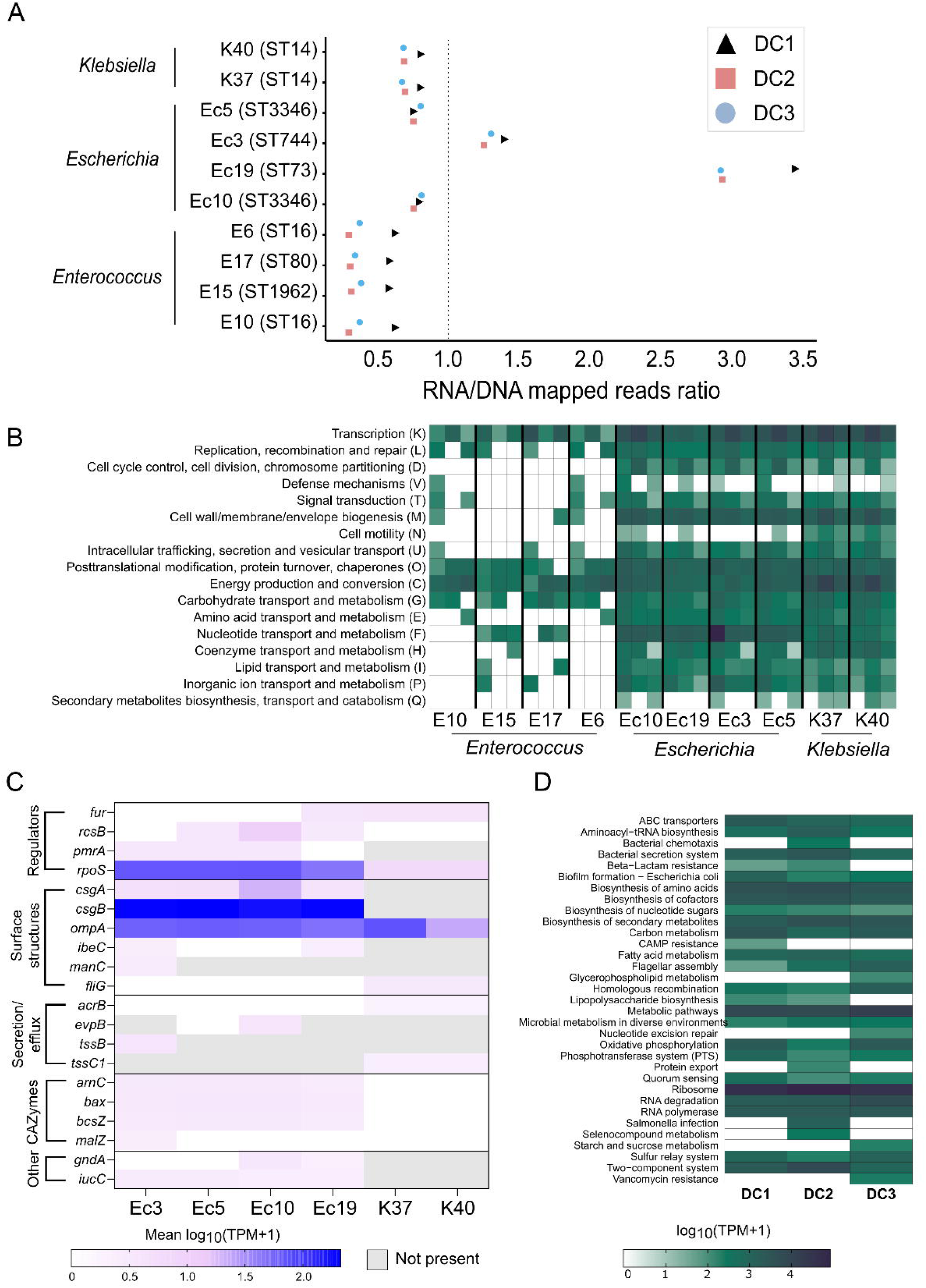
Metatranscriptomics analysis shows pathogen functional pathways and virulence genes are actively expressed *in situ*. A) RNA/DNA ratios of metagenomic and metatranscriptomic read mapping to isolates, indicating activity *in situ*. B) *In situ* transcriptional activity of each isolate in three independent water samples, shown as COG (Cluster of Orthologous Genes) category. C) average expression of Enterobacteriaceae isolate virulence factors and CAZymes *in situ* (mean log_10_(TPM+1) n=3). D) Ec19 KEGG pathways active *in situ*.

For a given pathogen, the term virulence factor refers to those genes that contribute to an overall virulent phenotype. The Enterobacteriaceae isolates also expressed virulence genes *in situ.* Those virulence genes that were expressed across the community (**Figure 1C**) also predominated on an isolate level, and encoded surface structures, transcriptional regulators, secretion systems and siderophores (**Figure 2C**). Virulence-associated regulators of gene expression (*fur*, *rcsB* and *pmrA*) were expressed in both *E. coli* and *K. aerogenes*, suggesting that genes within virulence regulons may be important for survival in freshwater. The master regulator of the general stress response, encoded by *rpoS*, was highly expressed indicating freshwater induced stress, likely due to nutrient limitation (47–49). RpoS also regulates the production of curli in *E. coli* (50). In all *E. coli* isolates investigated in this study, signature genes for curli production (*csgA*, *csgB*) were expressed in the freshwater, suggesting that this structure may facilitate survival *in situ*. Genes encoding a type 6 secretion system (*tssB*, *tssC1*) used for inter-species competition were also detected with strain specific expression profiles. A gene unique to and essential for flagella function, *fliG,* was active in *K. aerogenes* isolates indicating motility may be important for survival. Other cell surface structure genes were also expressed including *ompA*, which encodes the major outer membrane porin of *E. coli* (51), and *ibeC* and *manC*, both of which encode enzymes involved in synthesis of lipopolysaccharide.

### Investigation of a highly virulent environmental isolate of *E. coli* ST73

The *E. coli* isolate Ec19 was selected for further investigation given its high *in situ* transcription activity, and its membership of the global ST73 lineage that is associated with severe urinary tract infections (UTI) in many parts of the world including Australia (47). A core genome SNP-based phylogenetic tree showed Ec19 to cluster within a clade of ST73 isolates, including the prototypical uropathogenic *E. coli* (UPEC) strain CFT073 **(Figure 3A, Figure S4A)**. Ec19 harbours the AMR genes *bla*_CTX-M-24_ (an enzyme conferring resistance to third generation cephalosporins) and *tet(B)* (conferring resistance to tetracyclines), as well as a mutation in the topoisomerase *gyrA* (S83L) that confers fluoroquinolone resistance **(Figure 3A)**. Antibiotic susceptibility testing confirmed the predicted AMR phenotypes for Ec19 **(Figure 1E)**. Ec19 also possesses a wide range of UTI-associated virulence factors, including adhesins, toxins, immune modulation proteins, and iron acquisition systems **(Figure 3A)**. The virulence gene content of Ec19 is typical of the ST73 lineage, whose representatives typically possess a high number of virulence-associated genes in comparison to other common uropathogenic *E. coli* lineages (e.g., ST127, ST195, ST131, and ST1193) (52) **(Figure 3B)**. In the environmental metatranscriptomes, 12-16,000 reads mapped to the Ec19 genome (over 500 genes) at each of the three sampling dates; the most highly expressed included genes for house-keeping functions (e.g. transcription, translation, recombination), biosynthesis (lipopolysaccharide, amino acids, cofactors, nucleotides), catabolism (sugar and lipid catabolism, oxidative phosphorylation), transport (ABC transporters), and quorum sensing, whereas genes associated with chemotaxis, infection, and resistance to β-lactam antibiotics were variably expressed between water samples (**Figure 2D**). The synthesis of surface-associated virulence factors, specifically type 1 fimbriae and curli, was experimentally validated in the isolate using yeast agglutination and curcumin agar assays, respectively **(Figure 3C-D)**. As an independent means to validate curli production in Ec19, Western blotting was performed and showed that this strain produces curli , including at 37°C **(Figure S6B)**. Unlike the reference ST73 UPEC strain CFT073, Ec19 is cellulose-negative (**Figure 3E**) due to a transposon insertion in *bcsR* (**Figure S4B**). This surface factor profile, with production of curli but loss of cellulose production, has previously been linked to high virulence in various Enterobacteriaceae (53–56).

**Figure 3.**
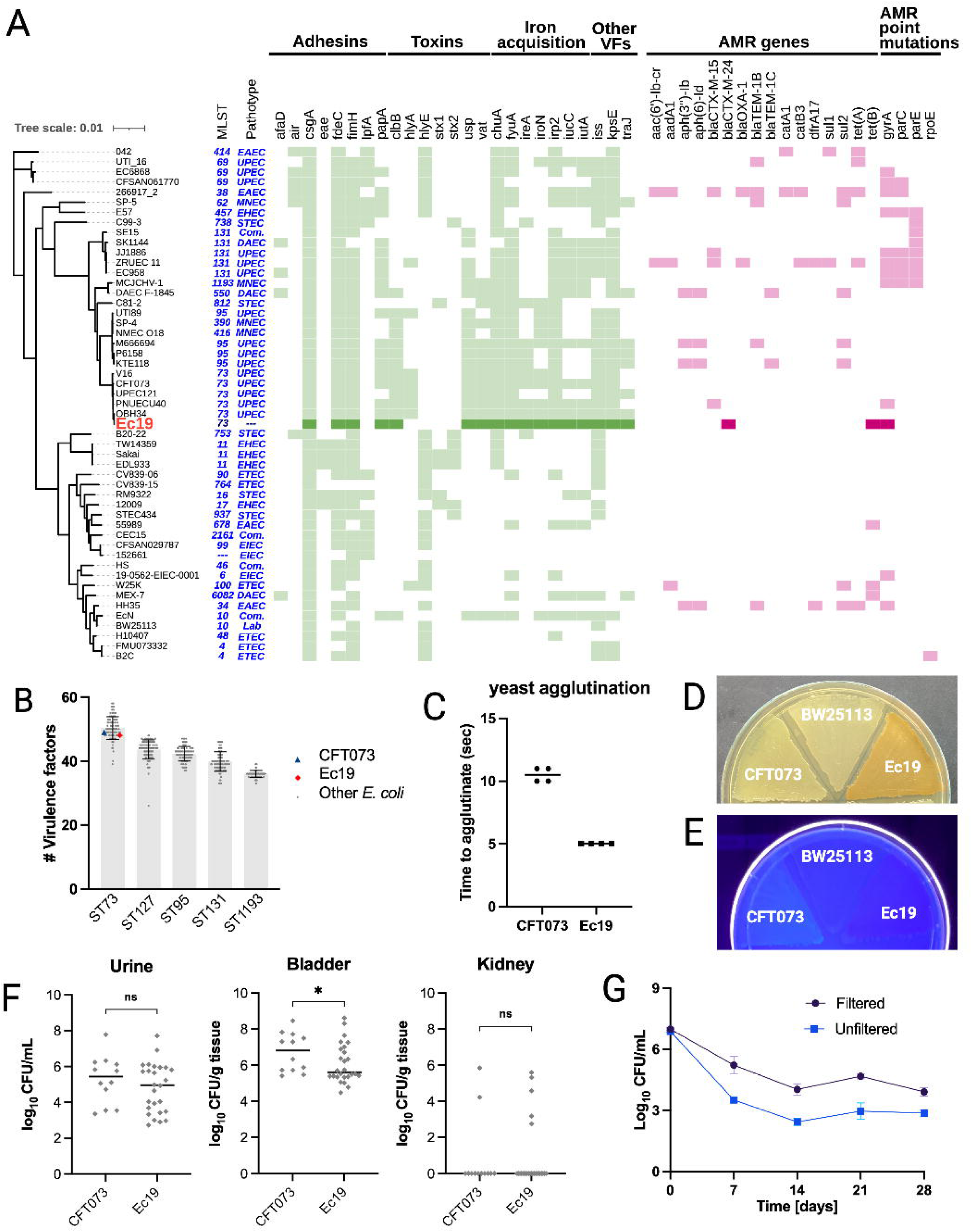
Virulence, AMR and environmental survival profile of Ec19. A) Phylogenetic tree comparing Ec19 and 50 pathogenic and commensal *E. coli* strains. The corresponding sequence type and pathotype of each strain are shown in blue. The presence-absence matrix of virulence and antibiotic resistance genes for each strain are shown in green and pink respectively, with the Ec19-associated genes shown as darker variants of those colours. B) The number of virulence factors predicted in Ec19, CFT073 and 100 randomly selected isolates from each of the five most prevalent UPEC lineages (ST73, ST127, ST95, ST131 and ST1193) from Enterobase. Error bars represent the standard deviation. C) Time to visible agglutination of CFT073 and Ec19 in the presence of yeast cells. Both strains exhibited strong agglutination capacity, reflective of type 1 fimbriae expression. D) Curli production of *E. coli* Ec19, CFT073 (curli-positive) and BW25113 (curli-negative) on YESCA agar supplemented with 18 µg/mL curcumin. Yellow and white colonies indicate positive and negative curli production, respectively. Darker yellow colonies correspond to higher curli production. E) Assessment of cellulose production in *E. coli* Ec19, CFT073 (cellulose-positive) and BW25113 (cellulose negative) on YESCA agar supplemented with 20 µg/mL calcofluor white. Cellulose producing isolates fluoresce under UV light. F) Mouse urinary tract colonisation following infection with *E. coli* CFT073 (n = 12) or Ec19 (n = 26). Viable bacteria in the urine, bladder and kidneys were enumerated 24 hours post-infection. The black horizontal line represents the median. Statistical significance was assessed using a Mann-Whitney test. ^ns^ = p > 0.05, * = p < 0.05. G) Survival of Ec19 in filtered or unfiltered creek water over 28 days.

Given it belongs to a major UPEC lineage and produces key colonisation factors, we assessed the capacity of Ec19 to cause UTI in a murine model. Ec19 was found to be virulent in mice and, compared to CFT073, showed similar colonisation of the mouse bladder and kidneys, as well as similar abundance in the urine **(Figure 3F)**. We next investigated the survival of Ec19 inoculated into water that was directly sampled from the Dandenong Creek (unfiltered), or in the same water filtered through a 0.22 µm Steritop filter, both incubated with consistent aeration. Whereas survival remained unchanged over five days in sterile water, in filtered water, Ec19 showed a rapid initial decline in viable cells followed by stabilisation over 28 days (**Figure S4C, Figure 3G**). The initial decline may have been partially mediated by phage-induced lysis, as plaque formation in Ec19 top agar co-incubated with creek water was observed following phage enrichment **(Figure S4D)**. Ec19 incubated with unfiltered creek water also showed an initial steep decline followed by stabilisation, with reduced survival over 28 days compared to filtered water **(Figure 3G)**. Note that microbial competition in the unfiltered water is likely to have declined over the 28-day incubation, as a separate experiment showed a reduction in recoverable microbes from the creek water over time **(Figure S4E)**. These results collectively showed that a virulent UPEC strain is capable of long-term survival, and that this survival is affected by the presence of the freshwater microbiota.

### Nitrogen and carbon metabolism, amino acid and nucleotide biosynthesis, and specific surface structures influence the survival of *E. coli* in urban water microcosms

We then performed transcriptomics and TraDIS in water microcosm assays to (i) identify how Ec19 reprograms its gene expression in creek water and (ii) identify genes required for survival. Since it was not possible to capture all environmental fluctuations of the natural sampling location, assays were performed at an environmentally permissive temperature of 25°C, which also represents the average water temperature of the Dandenong creek during summer (**Figure S1C**), and over short time frames (one day for transcriptomics, five days for transposon mutagenesis) to avoid confounding effects from a reduction in microbiome viability during storage (**Figure S4E**). While we cannot exclude batch effects, subsequent validation experiments showed high concordance between independent water samples (**Figure 5A**). In order to be comparable to published RNAseq datasets, transcriptomics analyses conducted in filtered creek water were compared to a rich media control grown at 37°C. This evaluation of the creek water environment revealed 536 differentially expressed genes, with 263 upregulated and 273 downregulated. It is not possible to distinguish which element of the environmental mileau these expression changes are in response to: altered nutrient conditions, temperature, osmolarity or other variables that differ between the test condition and the control (**Figure 4A**). Genes with increased expression were primarily involved in metabolic processes, particularly those for amino acids, nucleotides and essential nutrients such as nitrogen and phosphorus. Ribosomal proteins, bacterial secretion systems and ABC transporters facilitating amino acid and inorganic phosphate import also showed increased expression under creek water conditions (**Table S7**).

**Figure 4.**
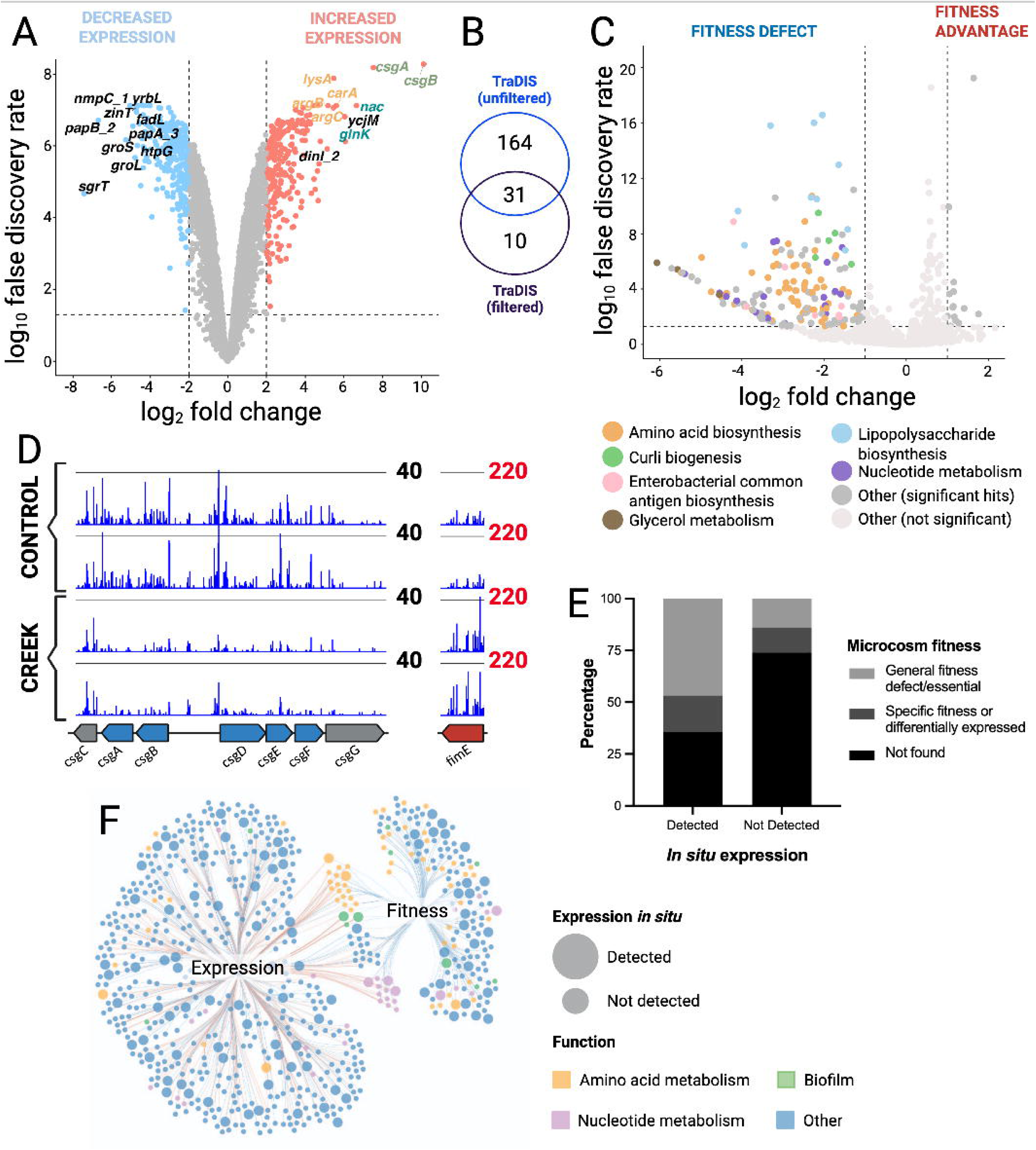
Mechanisms underpinning freshwater survival in Ec19. A) Microcosm RNA-seq volcano plot. Genes with higher expression are shown in red, lower expression in blue, and gene labels indicate those showing the largest changes in expression in creek water (ten highest and ten lowest) and are coloured based on function (yellow = amino acid biosynthesis, teal = nitrogen metabolism, green = curli, black = other). B) Venn diagram illustrating the number of microcosm TraDIS gene “hits” between unfiltered and filtered creek water conditions. C) Microcosm TraDIS volcano plot. Each gene is represented by a single plot. Refer to the legend below the plot for the corresponding biological function of each gene. D) Microcosm TraDIS insertion plots of key gene hits. The genes are represented by arrows at the bottom of the plot (grey = not a gene hit, blue = fitness defect hit, red = fitness advantage hit). E) Stacked bar chart illustrating the proportion of *in situ* expressed genes that are essential based on the TraDIS data F) Network map illustrating the intersection of the microcosm expression, *in situ* expression and fitness (unfiltered microcosm) datasets, produced in Cytoscape.

**Figure 5.**
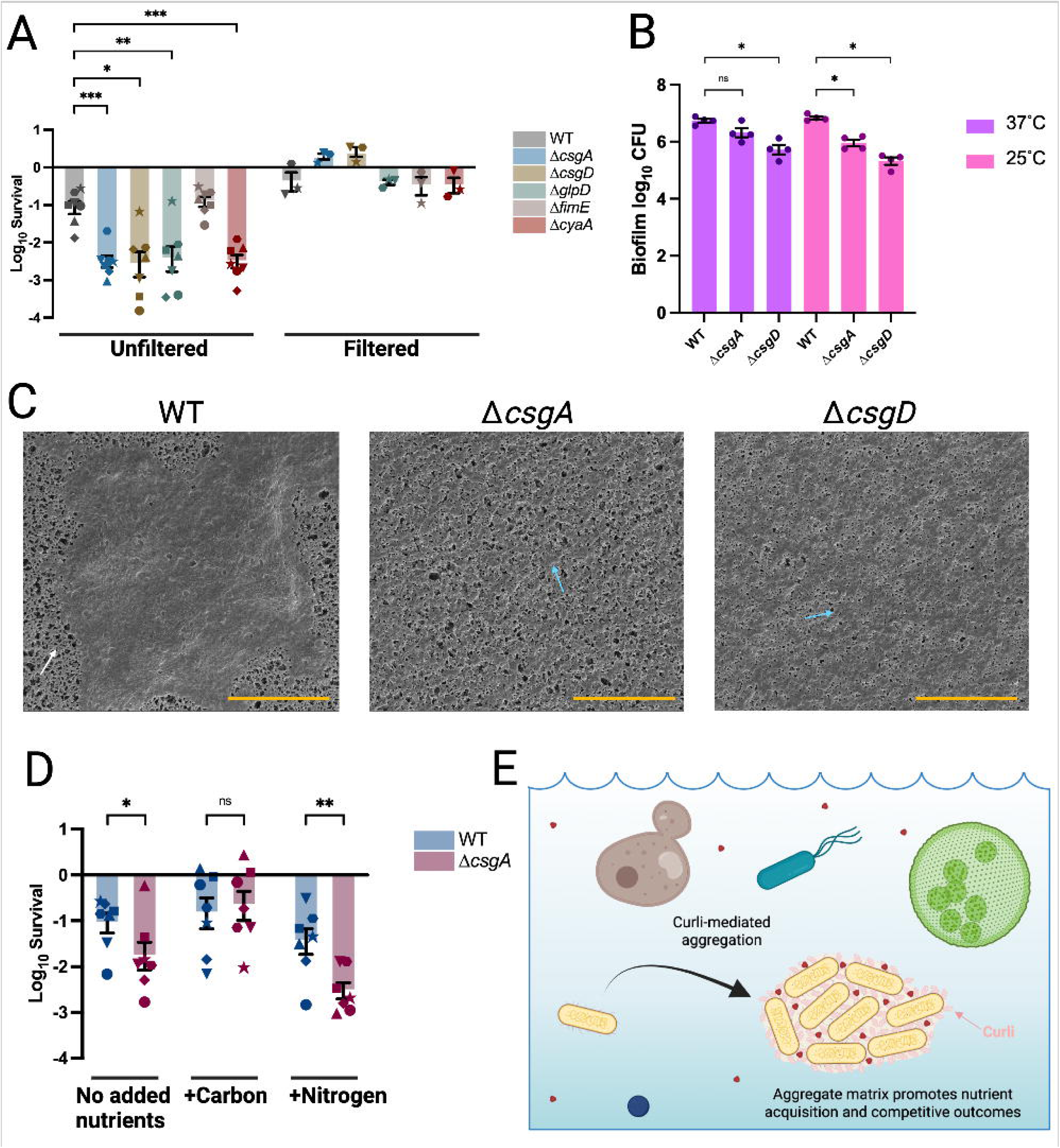
Comparisons between Ec19 WT and freshwater survival-implicated gene deletion mutants. A) Log_10_ survival of Ec19 WT and mutants after 5-day incubation in independent 1-day-old filtered or unfiltered creek water samples. B) Quantification of Ec19 WT and mutant biofilms grown at 37°C or 25°C for 24 hours. Biofilms were quantified based on CFU, and data points represent the average CFU from four technical replicates. C) Scanning electron microscopy of Ec19 WT and mutants following 5 days of creek water exposure at 25°C. Cells/aggregates were mounted on a multicellulose ester membrane which exhibits a mesh-like structure (white arrow). Blue arrows indicate individual cells. The orange scale bars represent 20 µm. D) Log_10_ survival of Ec19 WT and curli mutant after 5-day incubation in plain unfiltered creek water, or unfiltered creek water supplemented with either glucose as a carbon, or ammonium chloride as a nitrogen source. E) Schematic of model for curli-mediated aggregation effects *in situ*. Data points in A) and D) represent individual biological replicates, each measured in technical duplicate, and error bars represent SEM. Data points of the same shape indicate paired samples (i.e. biological replicates exposed to the same creek water batch). Statistical significance in A and D), and B was assessed using Mixed-effects model and RM Two-way ANOVA, respectively. ^ns^ = p > 0.05, * = p < 0.05, ** = p < 0.01, *** = p < 0.001.

Several metabolic pathways showed decreased expression, including those for degradation of amino and fatty acids, central carbon metabolism and sulphur metabolism, and various ABC transporters for sulphate, organic phosphate, galactose, and zinc acquisition (**Table S7**). Genes involved in amino acid biosynthesis (*lysA*, *argB*, *carA* and *argC*) and nitrogen metabolism (*nac* and *glnK*), respectively, were also highly induced (**Figure 4A**). The two most upregulated genes, *csgA* and *csgB*, encode the structural components of curli fibrils, a virulence-associated amyloid fibre that is extruded through the *E. coli* cell surface (**Figure 4A**) (57, 58). Lower temperatures, scarce nutrients, and low osmolarity are all known triggers to promote curli expression (57–60), which may explain the strong induction seen in creek water environment.

Genome-scale mutant fitness screening (TraDIS) was then used to interrogate the contributions of all non-essential Ec19 genes to survival in freshwater microcosms, and this analysis included both filtered and unfiltered water in order to capture microbiota effects on survival determinants. Testing the library under two different conditions yielded different results; the filtered freshwater condition resulted in 41 hits, whereas the unfiltered condition yielded 195 **(Figure 4B, Table S8, Table S9)**. The majority of hits in the filtered dataset were also detected in unfiltered conditions, suggesting that freshwater with its microbiota imposed more severe as well as distinct selective pressures on Ec19. We focussed on the unfiltered set for further analysis. In the unfiltered dataset, we identified several Ec19 metabolic processes, particularly for glycerol catabolism, and amino acid and nucleotide biosynthesis, that, when inactivated, resulted in a fitness defect under freshwater conditions **(Figure 4C)**. This was an expected consequence of the scarcity of nutrients in aquatic environments which necessitates that pathogens reprogram their metabolism in response (26). Changes to the Ec19 cell surface were also found to affect fitness, as inactivation of genes associated with biosynthesis of lipopolysaccharide, enterobacterial common antigen and curli (*csgA*, *csgB*, *csgD*, *csgE*, *csgF*), resulted in decreased fitness **(Figure 4C-D)**. Transposon insertions in *fimE*, which encodes a tyrosine-like recombinase that favours on-to-off phase switching of the promoter element that controls type 1 fimbriae expression, were associated with enhanced freshwater fitness, potentially due to elevated Type 1 fimbriae production which would be predicted to improve adhesion, aggregation and surface-associated biofilm formation (**Figure 4D**) (57, 60). There was a high degree of overlap (n = 49) between the genes with fitness effects (n = 205) and those that were differentially expressed in creek water (n = 460) that was greater than expected by chance (χ^2^ = 33.86, p < 0.0001).

Comparing the *in situ* and microcosm datasets revealed a high concordance between genes that showed high expression in the environment, and genes implicated as important through functional genomic approaches. Many Ec19 genes with detectable transcription *in situ* (214/450) were essential or near-essential based on the TraDIS results (defined as having logCPM of <3 in the control condition) (**Figure 4E**), and this enrichment was statistically significant (χ^2^ = 317.1, *p* < 0.0001). The 243 non-essential *in situ*-expressed genes were strongly enriched in the microcosm functional genomics datasets, with 80 of these showing freshwater-specific fitness impacts, differential expression, or both (χ^2^ = 62.55, *p* < 0.0001) (**Figure 4E**). Overall, 23% of Ec19 genes were implicated in freshwater survival through microcosm functional genomics experiments. However, considering only the subset of Ec19 genes that were expressed *in situ*, this proportion rose to 65%. Of the genes that were both required for fitness, and differentially expressed in the microcosm, these were enriched in functional categories corresponding to amino acid biosynthesis, nucleotide biosynthesis and biofilm formation (**Figure 4F, Figure S5**).

### Curli promotes aggregation to enhance nutrient competitiveness in freshwater microcosms

Based on the combined RNA-Seq and TraDIS datasets, we selected five creek water survival-implicated genes to further investigate, including *csgA* (encoding a structural subunit of curli fibrils), *csgD* (transcriptional activator of the *csg* operon), *glpD* (a representative gene of the glycerol metabolism pathway), *fimE* (encoding the type 1 fimbriae promoter switch recombinase) and *cyaA* (encoding adenylate cyclase). Targeted Ec19 deletion mutants were constructed by allelic exchange mutagenesis and their survival in freshwater microcosms evaluated under the same conditions as the microcosm TraDIS experiments, except that different batches of water from Dandenong Creek were used. Using seven independent one-day-old freshwater samples, we found that mutants of all fitness defect-associated genes (Δ*csgA*, Δ*csgD*, Δ*glpD* and Δ*cyaA*) had significantly lower survival compared to Ec19 (**Figure 5A**). The Δ*fimE* mutant did not exhibit the fitness increase that was observed for *fimE* transposon mutants in TraDIS (**Figure 5A**); it is possible that transposon insertion in *fimE* alters neighbouring gene expression while gene deletion does not, or that the phenotype observed by TraDIS only occurs in the context of a mixed population. Reduced survival phenotypes were only observed in unfiltered freshwater; filtering the water, which removes most bacteria (0.22 µm) but not necessarily viruses and dissolved factors (**Figure S4D-F**), resulted in similar or greater Ec19 mutant survival compared to the wild-type (**Figure 5A**). Considering only the three replicates where paired filtered and unfiltered water survival outcomes were available, Δ*csgA* and Δ*csgD* showed the greatest increase in survival following removal of the microbiota, with a log10 survival increase of 2.51 and 2.49, respectively, followed by Δ*cyaA* (1.9 log10 increase) and Δ*glpD* (1.49 log10 increase). The survival of the Δ*fimE* mutant did not change.

Finally, we investigated the physiological role of curli fibrils in the context of environmental survival. These fibrils form amyloid, and have been implicated in UPEC virulence (61) due to their proinflammatory properties and ability to bind fibronectin (62, 63) as well as being a component of *E. coli* biofilm extracellular matrix (57, 64, 65). We first tested the phenotypes of the Ec19 Δ*csgA* and Δ*csgD* mutants, validating their predicted inability to produce curli (**Figure S6A-B**), and their effect on static biofilm formation in rich media (**Figure 5B**). The Δ*csgA* mutant, which lacks the major structural fibrillar protein but still expresses CsgB, showed lower biofilm formation than Ec19 at 25°C. The Δ*csgD* mutant, which cannot activate expression of any of the curli fibrillar proteins, showed significantly lower biofilm formation at both temperatures (**Figure 5B**). Next, we tested whether curli fibrils provide protection from antimicrobial peptides, as protection from various harmful molecules has previously been reported (64, 66–68), and antimicrobial peptides can be produced by natural microbial communities (69). However, the Δ*csgA* and Δ*csgD* mutants of Ec19 did not show impaired survival in the presence of either polymyxin B or LL-37 at 25°C (**Figure S6A-D**).

We then investigated how the function of curli in biofilm may contribute to Ec19 survival in the freshwater ecosystem. Scanning electron microscopy of freshwater-incubated Ec19 was performed to determine how this surface factor may affect cellular arrangement *in situ*. Following incubation in a freshwater microcosm at 25°C, the curli-producing WT cells were densely packed in aggregates encased in a matrix, while cells of Δ*csgA* and Δ*csgD* appeared as a dispersed cells (**Figure 5C**). Finally, we sought to determine whether this curli-mediated aggregation provides benefits beyond protection and resistance. We hypothesised that aggregation and absorption to particulates may facilitate nutrient acquisition and help outcompete microbial competitors for resources. To test this hypothesis, we incubated Ec19 WT and Δ*csgA* in unfiltered creek water supplemented with a preferred carbon (glucose) or nitrogen source (ammonium chloride) to determine whether nutritional supplementation would reverse the survival defect of Δ*csgA*. Full complementation of the Δ*csgA* survival defect was observed upon carbon supplementation **(Figure 5D)**. Taken together, our data indicate that the extruded amyloid superstructure formed from curli fibrils promotes environmental survival of Ec19 by mediating aggregation, which subsequently facilitates carbon source acquisition to improve competitiveness in complex microbial communities.

## Discussion

Natural environments play a critical yet underexplored role in the transmission and evolution of opportunistic pathogens, a role that is likely to intensify as climate change and urbanisation increasingly perturb human–environment interfaces. Here, we integrated genome-resolved meta-omics with functional genomics to directly link *in situ* activity of pathogens to the genes required for their survival in a complex freshwater ecosystem. This approach addresses a major gap in One Health research by moving beyond detection of pathogens and resistance determinants, to identifying the mechanisms that enable environmental survival within native microbial communities.

Metagenomic and metatranscriptomic profiling revealed a freshwater community consistent with the characteristics of limnic systems, with Proteobacteria, Actinobacteriota, and Bacteroidota the most abundant phyla, and metabolic capabilities similar to other reported freshwater microbial communities (37, 70, 71). Although opportunistic pathogens occurred at low abundance, metatranscriptomics revealed that they were transcriptionally active in these communities, indicating that these bacteria are active members of the ecosystem. By mapping community RNA reads to isolate genomes from the *Escherichia*, *Klebsiella*, and *Enterococcus* genera, we demonstrate that genes in these isolates categorised as being associated with growth, metabolism, stress responses and virulence are expressed *in situ*. Ec19, a specific *E. coli* strain from a lineage capable of causing UTI, showed high transcriptional activity in the freshwater community. At the whole community level, various ARGs were also widespread and expressed, potentially reflecting endemic microbial competition and natural antimicrobial interactions in freshwater ecosystems, rather than solely anthropogenic selection (72, 73). To our knowledge, this is the first study to apply metatranscriptomics to interrogate the environmental physiology of opportunistic pathogens in a natural reservoir.

We demonstrated a capacity of Ec19 to survive for extended periods in freshwater, and also to cause infection in a mouse model of UTI. Functional genomics indicated that survival in freshwater depends on genes that fall into the broad functional categories of metabolic reprogramming, cell envelope maintenance, and cell surface appendages. Among these, curli biogenesis emerged as a central determinant of fitness: curli showed very high expression both in microcosms and *in situ*, and curli mutants had pronounced survival defects specifically in the presence of the native freshwater microbiota. Several of the additional fitness genes we identified such as *cyaA* are required for curli production (74, 75). That a survival advantage of curli was specifically observed in the presence of the freshwater microbiome suggests that these structures do not merely protect against abiotic stress but also enhance competitiveness within complex microbial communities.

Though curli have multiple reported roles, our data indicates a specific mechanism by which these structures promote freshwater survival (**Figure 5E**). Curli facilitate formation of non-attached aggregates (**Fig 5C**), a form of liquid-suspended biofilm found in aquatic environments (76–78). These meta-structures improve nutrient capture within this oligotrophic, competitive environmental niche and thereby increases the ability of Ec19 to compete with the native microbiota and survive (76, 79). Since these meta-structures would also entrap other strains and species, biofilm-associated survival in freshwater environments may also enhance opportunities for horizontal gene transfer, through bringing potential donors and recipients into proximity (80, 81). Our findings build on the known role of curli in *E. coli* biofilm formation (82, 83) to demonstrate for the first time that curli promote freshwater survival through nutritional competition.

*E. coli* ST73 is a globally disseminated pathogenic lineage that has previously been isolated from humans and multiple non-human sources including chickens with colibacillosis and household pets with UTIs (46, 84–91). Regardless of their source, most isolates from this lineage are phylogenetically clustered together, suggestive of zoonotic or environmental transmission capacity, not simply as human UPEC pathogens (87, 90, 92). We speculate that the presence of Ec19 in the creek is the result of contamination from humans or animals. While Dandenong Creek is a well-managed urban water system, its catchment area provides opportunity for contamination by wild and domestic animals, and potential for human sewage contamination due to waste containment failures such as septic tank failure or stormwater runoff. Indeed, there was a high transcriptional signal for Ec19 across all three sampling dates in our metatranscriptomics, indicating extended *in situ* survival of Ec19 or highly similar strains, frequent recontamination, or both. Ec19 was notable amongst our isolates for its high *in situ* activity, high virulence factor content and long-term freshwater environmental survival – a characteristic that was shown to be dependent on one of these virulence factors. This finding highlights that natural environments can act as important selective niches for clinically-relevant phenotypes in opportunistic pathogens; if confirmed to be general, this finding places even greater urgency on efforts to reduce environmental pathogen load.

More broadly, the approaches employed in this study can be widely applied to understand pathogen ecology and control. By providing a scalable framework that integrates environmental meta-omics with functional validation, this study offers a generalisable genome-scale methodology for dissecting the mechanisms by which pathogens persist in complex natural ecosystems. Such insights are essential for predicting and mitigating environmental contributions to the spread of infectious disease and antimicrobial resistance. Equivalent approaches can be applied to gain a genome-wide understanding of environmental adaptation of a range of pathogens, reveal how they respond to potential anthropogenic pressures such as warming, pollution, and antibiotic contamination, and understand how they respond to environmental engineering efforts to reduce their load.

## Materials and Methods

### Sample collection and bacterial isolation

Water samples were collected from Dandenong Creek at Tirhatuan Wetlands (−37.936845, 145.218046), a suburban site located on Bunurong Country in Victoria, Australia between (June 2023-March 2024) using sterile Schott bottles that were further disinfected with 80% v/v ethanol prior to sampling. All personnel wore appropriate PPE and disinfected nitrile gloves as above to minimise contamination. Bacterial isolation was conducted on samples from multiple dates from 7/6/2023-1/9/2023 including DC0 (10/08/2023), the isolation date of Ec19; isolation dates for all bacteria reported in this study are provided in **Table S3**. Metagenomics and metatranscriptomics was conducted on samples DC1 (22/11/2023), DC2 (15/02/2024), DC3 (11/03/2024). Additional samples from the same location were collected for microcosm experiments. Samples for metagenomics and bacterial isolation were stored at 4°C within 1 hour of sampling. Samples for metatranscriptomics were filtered immediately and filters were snap frozen with liquid nitrogen. Samples were vacuum filtered using 0.8 μm filter paper and 0.22 μm filter paper (Whatman®) in 200 mL aliquots to prevent blockage. Both 0.8 µm and 0.22 µm filter papers were plated directly onto CHROMagar™ Orientation (CHROM-O) or Slanetz and Bartley agar (SBA) then incubated aerobically at 37°C overnight. Burgundy colonies from CHROM-O plates were streaked onto MacConkey Agar and incubated overnight at 37°C to select for lactose-fermenting isolates, which were denoted as presumptive *Escherichia* spp. Blue colonies on CHROM-O plates were re-streaked once on CHROM-O plates to obtain single colonies, which were then streaked onto Simmons Citrate Agar with Inositol (SCAi) and incubated for 48 hours at 37°C to select for yellow colonies, which were denoted as presumptive *Klebsiella* isolates. Maroon colonies from SBA were streaked onto Bile Esculin Agar (BEA) and incubated overnight at 37°C to select for brown colonies, which were denoted as presumptive *Enterococcus*. Well-isolated colonies for each genus were picked and streaked onto nutrient agar prior to 16S rRNA sequencing. All bacteria isolated or used during this study are listed in **Table S3, Table S4** and **Table S10.**

### 16S rRNA gene sequencing

Isolated colonies for each isolate were resuspended in water, boiled at 100°C for 10 minutes before centrifugation at 13,000 rpm. 5 µL of supernatant was added to a PCR master mix containing 1 µL of 10 µM 1492R reverse primer (5’-GGTTACCTTGTTACGACTT-3’), 1 µL of 10 µM 27F forward primer (5’-AGAGTTTGATCCTGGCTCAG-3’), 25 µL *Taq* 2X Master Mix (New England Biolabs), and 18 µL nuclease-free water. Samples were subjected to thermocycling, 30s denaturation at 95°C followed by 30s annealing at 50°C and 30s extension at 68°C for 30 cycles with a final extension at 68°C for 5 minutes. PCR products were excised from 1% agarose gel and the DNA cleaned using a Monarch DNA Gel Extraction Kit (New England Biolabs). Samples were sent for Sanger sequencing at Micromon Genomics (Monash University, Clayton, Australia). Preliminary genus identification based on the 16S rRNA sequence was performed using BLASTn (88).

### Isolate whole genome sequencing

Isolate genomic DNA (gDNA) extraction was performed using the Monarch Genomic DNA Purification Kit (New England Biolabs) as per the manufacturer’s instructions. gDNA was quantified using Qubit (Thermo Fisher Scientific) and sent for whole genome sequencing 150 bp paired end Illumina sequencing with Azenta. Additionally, Ec19 gDNA was sent for long read sequencing (Oxford Nanopore, Alfred Hospital). Hybrid assembly of Ec19 was completed using Unicycler v0.050+galaxy1 (Galaxy Australia) and short-read assemblies of other isolates were completed with Unicycler v0.5.1 (93). Isolates were typed using pubMLST (40) or Kleborate (39). The Ec19 genome was annotated with Prokka (v1.13.6+galaxy1) using a Genus-specific BLAST database for *E. coli* (−-usegenus –genus “escherichia” –species “coli), and with search for non-coding RNAs enabled (−-rfam) (94).

### Metagenomics

Single 0.22 um filters were used for DNA extraction using DNeasy PowerSoil Pro kit (QIAgen) as per the manufacturer’s instructions with 30 μl elution volume. Three extractions were completed per independent biological sample (DC1, DC2, DC3). Additionally, three technical replicates for each biological replicate were sent for Illumina 150 bp paired-end sequencing at (BMKgene, Beijing) to generate 15G raw data and average ∼65 million reads per sample. Adaptor sequences were trimmed using BBduk from the BBtools package (BBMap - Bushnell B. - sourceforge.net/projects/bbmap/), before assessing read quality using FastQC (v0.12.1) (95). PhyloFlash (v3.4.2) (96) was used to taxonomically profile the community with reconstruction of SSU rRNA genes from short-read shotgun metagenomic data. Forward reads were screened for functional genes using DIAMOND BLASTx (97) against an in-house database of 52 metabolic marker genes (98), which indicate the presence of key metabolic pathways involved in energy production and conservation. Hits were normalised to ribosomal genes to produce copies per genome (cpg) as previously described (98). Short reads were also screened with metaVF (38) and resistance gene identifier (RGI) (99) to quantify potential virulence factors and antimicrobial resistance genes in the community. Reads per kilobase million (RPKM) was calculated for each metaVF hit, which were then grouped by virulence factor category. Similarly, RGI RPKM was calculated for RGI hits, which were summed by drug class. Short reads were also mapped to isolate genomes using the subread package (subread-align v2.0.8) (100).

### Metatranscriptomics

Seven 0.22 μm filters were used for total RNA extraction from each biological sample using RNeasy PowerSoil total RNA kit (QIAgen) with the following modifications: one 0.22 μm filter was mixed with 357 μl PowerBead Solution, 35.7 μl Solution SR 1, 114 μl Solution IRS, and 500 μl phenol/chloroform/isoamyl alcohol (Merck) in a Lysing Matrix E 2-ml tube (MP Biomedicals), and homogenised at 6 m/s for 40 s (FastPrep 24, MP Biomedicals), then the tubes were centrifuged at 16,000 x g for 10 min before pooling the aqueous phase of seven tubes into a clean 15 ml Collection Tube, where all additional steps were conducted as per manufacturer’s instructions with 30 μl elution volume. The extracted RNA solution was treated with TURBO DNA-free™ Kit (Thermo Fisher) to reduce the DNA contamination. Total community RNA was sequenced at the Hudson Institute of Medical Research Genomics Facility (Melbourne, Australia). Libraries were prepared with Illumina Stranded Total RNA with RiboZero Plus treatment and sequenced using NovaSeq 6000 SP to generate 100 bp paired end reads (∼43 million reads per sample). Metatranscriptomics data were profiled with PhyloFlash and screened for metabolic marker genes as described previously (98, 101), although prior to screening, RiboDetector was used to computationally deplete ribosomal RNA (102)(98), and hits were normalised to RPKM.

Metatranscriptomics reads were mapped to isolate genomes to determine which isolates were active *in situ*, and understand which genes they use to survive. To reduce reporting of cross mapped reads, short-read metagenomic data was filtered using Kraken2 (99) for Enterobacteriaceae or Enterococcaceae reads, which were subsequently assembled into contigs using Metaspades (100). Enterobacteriaceae or Enterococcaceae contigs were used to build a subread index with individual isolate genomes from the same family (e.g. Ec19 + Enterobacteriaceae contigs used to build subread index). Metatranscriptomics reads were mapped with the subread package (subread-align v2.0.8) to isolate genomes in the context of metagenome reads from the same family, where only uniquely mapping reads with less than three mismatches were retained (97% identity and multimapping reads were removed from the analysis). Samtools (v1.21) was used to sort and index bam files (103), mapped reads were quantified using featureCounts and normalised to transcripts per million (TPM) relative to mapped reads (100, 104). Genes that had mapped reads were annotated using eggnog-mapper v2 (v2.1.12 http://eggnog-mapper.embl.de/) (105) and collapsed into Clusters of Orthologous Genes (COGs), KEGG pathways, KEGG modules or CAZyme categories. Abricate (v1.4.0) (Seemann T, Abricate, Github https://github.com/tseemann/abricate) was used to annotate *in situ* expressed virulence genes in isolate genomes. Additionally, metatranscriptomics datasets were mapped to the Ec19 genome and output forward reads were screened for pathways using DIAMOND BLASTx (97) against a local KEGG database. Short reads were also screened for virulence factors and ARGs with metaVF and RGI as described above.

### Antibiotic susceptibility testing

Sensititre Plates (Thermo Fisher Scientific) were used to determine the antibiotic susceptibility profiles of the 10 isolates that underwent WGS. For the gram-negative isolates, including *Klebsiella* and *E. coli,* the Sensititre Gram Negative MIC GNX2F Plate (Thermo Fisher Scientific) was used. For the gram-positive *Enterococcus* isolates, the Sensititre™ Gram Positive GPALL3F AST Plate (Thermo Fisher) was used. Three to five colonies of each isolate were picked from plates using a wire loop and suspended in 4 mL of PBS. Optical density was adjusted via dilution or further inoculation to an OD_600_ of 0.15-0.18, equivalent to 0.5 McFarland Standard. For the non-fastidious gram-negative isolates, 10 µL of cell suspension was added to 11 mL Mueller-Hinton Broth (MHB). For gram-positive isolates, 30 µL was added to 11 mL MHB. Each well in the 96-well Sensititre plates was inoculated with 50 µL of the MHB suspension before they were sealed and incubated overnight at 37°C. Wells for positive and negative growth controls were included on the plate. Each well was evaluated for growth after incubation.

### Phylogenetic analysis of Ec19

Phylogenetic analyses were performed on Ec19 and 50 pathogenic and commensal *E. coli* strains randomly retrieved from NCBI. Genome FASTA files of each strain were used as inputs in Prokka (v1.14.6) to generate a gff3 file for core genome alignment in Roary (v3.13.0). Poorly aligned sequences from the genome alignment file were trimmed off using trimAI (v1.5.0) before being used as input for iqtree (v2.3.6) for phylogenetic tree construction. The tree was visualised using iTOL v6. Virulence genes were identified using the web version of VirulenceFinder (v2.0.5) (106) and ResFinder (v4.5.0) (107). For comparisons of virulence gene content between lineages, Abricate (https://github.com/tseemann/abricate) was used with a curated *E. coli* virulence factor database (108) on 100 strains from ST127, ST195, ST131, and ST1193 (105)

### Yeast agglutination assay

The capacity of bacteria to express D-mannose-binding phenotype was assayed by their ability to agglutinate yeast (Saccharomyces cerevisiae) cells on glass slides as previously described (108). Aliquots of bacterial suspensions at OD_550_ = 0.5 and 5% yeast cells were mixed, and the time to agglutination was recorded. D-mannose-binding specificity was determined by performing the assay in the absence and presence of 50 mM methyl a-D-mannopyranoside (108).

### Curcumin agar assay

Curli production was assessed based on binding to the dye curcumin (109). Ec19 WT and curli mutants (Δ*csgA* and Δ*csgD*) were streaked onto YESCA agar plates supplemented with 18 µg/mL curcumin. The plates were wrapped in foil and incubated at 25°C for 48 hours. Curli-positive and -negative *E. coli* were identified as yellow and white colonies, respectively.

### Calcofluor white assay

Cellulose production was assessed based on the strong binding of a UV-fluorescent stain, calcofluor white, to cellulose (53, 109). Ec19 WT was streaked onto YESCA agar plate supplemented with 20 µg/mL calcofluor white. The plates were wrapped in foil and incubated at 25°C for 48 hours. After incubation, the plates were exposed to a UV transilluminator to assess calcofluor white binding. Cellulose-producing *E. coli* were identified as fluorescent white colonies under UV. *E. coli* CFT073 and BW25113 were used as cellulose-positive and negative controls, respectively.

### Mouse infection experiments

The C57BL/6 female mouse model of UTI was employed as previously described (PMID: 38383596, 30131362). Bacteria were grown statically at 28°C for 24 hours in LB no salt. Bacteria (∼5 × 10^8^ CFU in 20 μl PBS) were injected transurethrally into female C57BL/6 mice (8–10 weeks). Bacterial loads corresponding to each strain in the urine, bladder, and kidney at 24 hours postinfection (hpi) were enumerated by plating onto LB agar. Statistical analyses were performed using the two-tailed Mann-Whitney test (Prism10; GraphPad). All work involving mice was approved by the University of Queensland Animal Ethics Committee (AEC approval number 2022/AE000469) (53, 74).

### Microcosm survival assays

Overnight cultures of Ec19 were grown in 5 mL LB at 37°C. The cultures were centrifuged at 8000 rpm for two minutes to pellet the bacterial cells, then resuspended in 5 mL M63 minimal medium without a carbon source (M63-NC) to an OD600 of 0.8. 1 mL of normalised suspension was added to a sterile conical flask containing 49 mL unfiltered freshwater from Dandenong creek, or water that had been filtered through a Millipore^Ⓡ^ Steritop^Ⓡ^ 0.22 µm vacuum filter. The flask was incubated at 25°C with shaking at 100 rpm for 28 days. An aliquot of the water sample from each flask was collected for 10-fold serial dilutions and CFU measurements at specified time points, using tetracycline-supplemented agar (10 μg/ml) to allow growth of Ec19 which preventing growth of other microbes. Freshwater survival assays for validation of TraDIS experiments were performed in the same way as the TraDIS incubations.

### Microcosm RNA-Seq

Ec19 cells were prepared as above and incubated in filtered freshwater at 25°C with 100 rpm shaking for 24 hours. After incubation, the Ec19 cells in the water samples were collected by centrifugation and flash-frozen at -80°C within 10 minutes. RNA was purified by phenol-chloroform extraction as previously described (110). Purified RNA samples were sent to the Hudson Genomics Facility in Clayton, Victoria, Australia, for library construction and Illumina sequencing. Input overnight cultures of Ec19, prepared in LB at 37°C, were used as the rich media control condition. Experiments were performed in biological triplicate.

### RNA-Seq analysis

RNA-Seq analysis was performed using software implemented in Galaxy Australia. Paired-end sequencing reads were assessed for quality using FastQC (Galaxy v0.74+galaxy1). The sequencing reads were mapped against the *E. coli* Ec19 reference genome using BWA-MEM (Galaxy v0.7.18). Mapped sequencing reads for each Ec19 genome feature were tallied using featureCounts (Galaxy v2.0.3+galaxy2), using the Prokka-annotated Ec19 genome gff3 file (−t CDS, -g ID). FeatureCounts tables generated across three control and three water-treated replicates were compiled into a single count matrix using the Generate count matrix (Galaxy v1.0) software. The count matrix table was uploaded as an input to Degust (https://degust.erc.monash.edu/) for differential expression analysis (111). Differentially expressed genes were identified using a threshold of log_2_FC ≥ ∓ 2 and false discovery rate (FDR) < 0.05. KEGG pathway enrichment analysis was performed to identify biological pathways that were differentially expressed. Differentially expressed Ec19 genes were first assigned to *E. coli* CFT073 orthologs using reciprocal BLAST. The CFT073 locus tags were then converted into UniProt IDs using the UniProt ID mapping service (https://www.uniprot.org/uploadlists/) using the *E. coli* CFT073 [199310] database. The converted UniProt IDs were used as inputs for the KEGG Orthology Based Annotation System (KOBAS) web server (://bioinfo.org/kobas/genelist/) using the *E. coli* CFT073 database and Fisher’s exact test with Benjamini and Hochberg (1995) FDR correction for statistical significance (112).

### TraDIS mutant library construction

The *E. coli* Ec19 TraDIS mutant library was constructed using the transposon delivery vector pDS1028 (**Table S11**), a suicide Tn delivery vector containing a miniTn5-derived transposon conferring chloramphenicol resistance (113). Ec19 was conjugated with pDS1028-carrying *E. coli* JKE201 (a diaminopimelic acid (DAP)-dependent biparental mating strain (114)) in a 2:1 donor-recipient ratio. The Ec19-JKE201 conjugation mix was spotted on LB agar supplemented with 150 µM DAP for 1 hour. The conjugation patch was collected using a sterile swab and resuspended in 25% glycerol. Aliquots of the conjugation mix were spread onto 27x27 cm bioassay plates containing LB agar + 35 µg/mL chloramphenicol to select for transposon insertion mutants, at a density of ∼10,000 colonies per plate. A total of 40 bioassay plates were used, to obtain up to 300,000 transposon mutant colonies, Transposon mutants were harvested, resuspended in 25% glycerol in 0.5x PBS, and mixed at 25°C and 200 rpm for 30 minutes to evenly disperse mutants. The pooled transposon mutant library was stored at -80°C until future use.

### Microcosm TraDIS

Cultures of the Ec19 TraDIS library were grown in 5 mL LB at 37°C, 100 rpm overnight, and washed and normalised as for the RNA-seq experiments. Filtered and unfiltered freshwater conditions were used. For the filtered condition, 1 mL of normalised TraDIS library suspension was added to 49 mL fresh creek water (collected one day before the experiment) that had been filtered using the Millipore® Steritop® Vacuum Bottle Top Filter (0.22 µm pore size). For the unfiltered condition, 1 mL of normalised TraDIS library suspension was transferred into pre-washed SnakeSkin™ dialysis tubing (3.5 kDa MWCO), with its ends tied to contain the suspension, and placed in a sterile conical flask containing 49 mL fresh unfiltered water. Dialysis tubing was necessary in unfiltered water to prevent phage infection of the Ec19 library and to ensure enrichment of Ec19 rather than the autochthonous Dandenong creek water microbiota, but was deemed unnecessary in filtered water as the higher Ec19 survival allowed maintenance of sufficient library diversity without excessive population structure distortion by phage infection (115). Flasks from both experimental conditions were incubated at 25°C, 100 rpm for five days, at which point Ec19 cells were harvested by centrifugation, resuspended in 100 µL sterile deionised water, and 10 µL of this suspension was outgrown overnight in 5 mL LB at 37°C, 100 rpm. Control libraries were overnight cultures of the Ec19 library that were centrifuged, resuspended and outgrown as for the experimental samples. Genomic DNA was extracted using the DNeasy® Blood & Tissue kit (Qiagen). DNA samples were submitted to the Ramaciotti Centre for Genomics (University of New South Wales) for TraDIS library construction and sequencing using a previously described library construction protocol (116) and sequencing primers previously described for pDS2018 (117). The control, unfiltered and filtered water conditions were performed in biological triplicate.

### TraDIS analysis

FastQ files from Ramaciotti were analysed using the Bio::TraDIS pipeline (116) with modifications. Read were assessed for quality using FastQC (Galaxy v0.12.1), and the poor-quality and illumina adapter sequences were trimmed using Trimmomatic (Galaxy v0.36.6; default parameters with the ILLUMINACLIP step included). The trimmed reads were mapped against the Ec19 genome using the *bacteria_tradis* command of the Bio::TraDIS pipeline, using Smalt mapper with a 95% sequence identity threshold (−-smalt_y 0.95) and the pDS1028 tag sequence (“TAAGAGACAG”) with two mismatches allowed. The resulting plot files were assigned to genomic features of Ec19 using the *tradis_gene_insertion_sites* function (default parameters) and the resulting gene-wise read count values used to generate a count matrix. Control and freshwater treatment count matrices were uploaded as inputs to Degust for differential expression analysis (https://degust.erc.monash.edu/) to identify genes with differentially abundant transposon insertions. Differentially abundant genes were defined as those with at least 10 transposon insertions in at least one replicate of either the control or experimental condition, and meeting a threshold of log_2_FC ≥ ∓ 1 and a false discovery rate (FDR) < 0.05.

### Mutagenesis of *E. coli* Ec19

Single gene deletion mutants were constructed in Ec19 using an allelic exchange mutagenesis method optimised for multidrug-resistant gram-negative bacteria (114), with modifications. The flanking regions upstream and downstream of the target gene were PCR-amplified using Q5 High-Fidelity 2X Master Mix (New England Biolabs), and the two fragments joined and cloned into pFOK (linearised with BamHI-HF®) by Gibson assembly using the NEBuilder® HiFi DNA Assembly Master Mix. Reaction contents were transformed into chemically competent *E. coli* JKE201 and selected on LB supplemented with 100 µg/mL ampicillin and 150 µM DAP. Mutagenesis vectors were transferred to *E. coli* Ec19 via conjugation. Donor cells (grown on LB agar at room temperature overnight) and recipient cells (grown on LB agar + 100 µg/mL ampicillin and 150 µM DAP at 37°C overnight) were resuspended in sterile PBS and adjusted to cell densities of OD_600_ = 40 and OD_600_ = 20, respectively. 50 µL of a 1:1 mixture of donor and recipient was spotted to an LB agar plate supplemented with 150 µL DAP, and incubated for 2 hours at 37°C. Conjugation patches were collected and resuspended in 1 mL sterile PBS using a sterile swab, and the suspension streaked on LB agar + 50 µg/mL Kanamycin to select for transconjugants which were confirmed by colony PCR using FS199 and FS205 primers. Single-crossover Ec19 mutants were streaked on LB agar + 20% w/v sucrose and 0.5 µg/mL anhydrotetracycline and incubated at room temperature for 48 hours to select for putative double-crossover Ec19 colonies. Putative double-crossover Ec19 colonies were patched on LB agar + 20% w/v sucrose and 0.5 µg/mL anhydrotetracycline and LB agar + 50 µg/mL Kanamycin to confirm loss of the vector. Both plates were incubated at 37°C overnight. Verified double-crossovers were screened for the loss of target gene via colony PCR. All primers used in this study are listed in **Table S12**. Genomic DNA from mutants was extracted using the QIAGEN DNeasy Blood & Tissue kit and Illumina sequenced by Azenta. Sequencing reads were mapped to the EC19 reference genome to confirm the deletions, and analysed using Snippy (v4.6.0+galaxy0) to check for unrelated secondary mutations.

### SDS-PAGE and Western blotting of curli

5 µL of overnight culture were spotted onto YESCA agar for incubation at either 25°C, 30°C or 37°C for 24 hours, and the growth culture were collected using sterile cotton swabs and resuspended in 1 mL PBS. The OD_600_ of the suspension were normalised to 1 before they were centrifuged to pellet bacterial cells and discard the supernatant. The cell pellets were resuspended in 40 µL formic acid, followed by drying of the suspension using the SpeedVac at 65°C for 40 minutes. The dried cell lysate were used for SDS-PAGE, and the electrophoresis was performed at 160 V, 400 mA for 70 minutes. Proteins in the gel were semi-dry transferred onto a polyvinylidene difluoride membrane using the Biorad Tran-Blot Turbo System at 1.3 A, 25 V for 7 minutes. Membranes were blocked in 1% skim milk and 1% bovine serum albumin in Tris-buffered saline with 0.1% Tween-20 (TBS-T) at 4°C overnight before incubation with rabbit anti-CsgA antibody (1:2000 dilution in 1% skim milk and 1 % BSA in TBS-T) or rabbit anti-OmpA antibody (1:10000 dilution in 1% skim milk and 1 % BSA in TBS-T) for 1 hour. The membranes were washed three times, five minutes each with TBS-T before incubation with a horseradish peroxidase-conjugated goat anti-rabbit antibody for 1 hour. The membranes were washed three times, five minutes each with TBS-T before chemiluminescent detection of protein using ChemiDoc MP imaging system (Biorad).

### Validation of mutant survival phenotypes

Survival of isogenic Ec19 mutants was evaluated using the same sample preparation, dialysis tubing methodology and incubation conditions as the Microcosm TraDIS described with the following modifications: the inoculum was a pure culture (either an Ec19 mutant or WT control), and recovered cells were vortexed at 2500 rpm for 2 minutes, followed by sonication for 10 minutes using a sonication bath in order to break up aggregates and allow accurate enumeration. The Ec19 suspension was then 10-fold serially diluted for CFU measurements (T_5_). A before-exposure control (T_0_) was included by measuring the CFU of the normalised Ec19 suspension. Log_10_ survival of each Ec19 strain was calculated using the formula T_5_/T_0_.

### Nutrient supplementation survival assay

This assay is identical to the mutant validation experiments above, but the water samples were supplemented with 0.4% w/v glucose or 9.5 mM ammonium chloride as carbon and nitrogen source, respectively.

### Biofilm assay

Biofilms were grown by diluting overnight cultures of WT or mutant Ec19 1:100 in Mueller-Hinton broth, then incubating 100 µL aliquots of this suspension in a 96-well plate (Nunc™ MicroWell™ 96-Well, Nunclon Delta-Treated; Thermofisher Scientific) at 37°C or 25°C, 75 rpm shaking for 24 hours. After incubation, the liquid suspension in the wells was removed and the surface-attached biofilms were washed twice in 200 µL sterile PBS. Biofilms were quantified by collecting the biofilm using a sterile cotton swab soaked in 100 µL sterile PBS. The cotton swab containing the detached biofilms was suspended in a 1.5 mL tube containing 900 µL sterile PBS with the cotton tip broken off and placed inside the tube. The tubes were vortexed at 2500 rpm for 2 minutes, then sonicated at max intensity for 10 minutes. The bacterial suspension inside the tubes were sampled for 10-fold serial dilutions and CFU plating.

### Scanning electron microscopy

Suspensions of freshwater-exposed cells were prepared using the same dialysis tubing methodology and incubation conditions as the Validation of mutant survival phenotypes assay above. The suspensions were recovered from their dialysis tubing and a 50 µL aliquot of this were spotted onto a multicellulose ester membrane (filter type 0.22 µm, MF-Millipore®) before leaving it to dry. Afterwards, the water-exposed cells were fixed with 200 µL 2.5% glutaraldehyde in 0.1M cacodylate buffer and left to incubate at 4°C overnight. After incubation, excess glutaraldehyde was removed by placing the membrane on top of a paper towel, thereby absorbing the fixative, followed by washing the membrane twice with 200 µL 0.1M Cacodylate buffer for 5 minutes, with removal of the buffer in-between by placing the membrane in a paper towel. The bacterial cells on the membrane were then dehydrated with multiple cycles of ethanol treatment at increasing concentrations: 50%, 75%, 90% and absolute ethanol. For each cycle of treatment, 200 µL of ethanol was added onto the membrane to dehydrate for 10 minutes, followed by placing the membrane on top of a paper towel to remove excess ethanol. After ethanol treatment, the membranes were left to air dry at 37°C overnight. After drying, the membranes were placed on an SEM metal stub containing carbon adhesive tabs before sputter-coating them in 20 nm of gold using the Leica EM ACE600 Sputter coater. The gold-coated samples were imaged using the FEI Nova NanoSEM.

### Statistics

Statistical analyses were performed using the Mixed effects analysis with Holm-Šídák’s multiple comparisons test, two-way RM ANOVA with Dunnett’s comparisons test or Chi-squared test as part of GraphPad Prism 10.

## Acknowledgments

We thank Professor Perran Cook and members of the Short, Greening and Peleg research groups for helpful discussions. We acknowledge that this project was conducted on the traditional homelands of the Bunurong people, and recognize the Bunurong people as the sovereign First People of their Country.

## Author Contributions

Conceptualisation F.S, C.G, T.W, C.N, T.L

Methodology T.W, T.J, J.S, C.N, L. V-A, F.S, C.G, M.A.S, T.N., Y.Q.

Software Y.L, W.S, T.W, C.N Validation C.N, T.W, F.S, C.G

Formal analysis T.J, J.S, C.N, T.W, N.T.K.N, T.N., Y.Q.

Investigation T.J, J.S, T.N.TN, G.H, C.N, T.W, N.T.K.N, T.N., Y.Q.

Resources C.G, F.S, Y.L, W.S, M.S, S.H.

Data curation T.W, C.N, F.S, T.J, J.S

Writing - Original Draft T.W, C.N, F.S, C.G

Writing - Review & Editing All authors

Visualisation T.W, C.N, F. S.

Supervision F.S, T.W, G.H, C.G, M.A.S

Project administration T.W, F.S

Funding acquisition C.G, F.S, T.L

## Competing Interest Statement

The authors declare that no competing interests exist.

## Classification

Biological Sciences; Microbiology

## Notes

### Competing Interest Statement

The authors have declared no competing interest.

### Summary of Updates

The manuscript has been modified to include additional data, revised figures and additional intepretation incorporated after the peer review process.

### Funding

- National Health and Medical Research Council: APP1178715, 2037698, APP2016330
- Australian Research Council: FT240100502, DE200101524, DP230101930

