## Supplementary figures S1-S6 and supplemental note for "Linking *in situ* activity to fitness reveals that clinically-relevant factors underpin pathogen survival in environmental reservoirs"

**This supplementary appendix contains:**

**Supplementary Tables – in *Supplementary_tables_S1-S12.xlsx***

**Table S1 Community ARG summary**

**Table S2 Community virulence factor summary**

**Table S3 Isolate summary**

**Table S4 Confirmed isolates**

**Table S5 Mapped metaT summary**

**Table S6 Mapping statistics and RNA/DNA ratio**

**Table S7 RNAseq KEGG pathways**

**Table S8 TraDIS summary statistics**

**Table S9 RNAseq TraDIS data**

**Table S10 Strains used for molecular microbiology**

**Table S11 Plasmids used**

**Table S12 Primers used**

**Supplementary Figures**

**Figure S1 Sampling sites and freshwater microbial community and target genera abundance and expression**

**Figure S2. Metabolic gene abundance and expression in freshwater microbial communities and isolates**

**Figure S3. COG categories and KEGG modules actively expressed by isolates *in situ***

**Figure S4 Creek water survival, microbiome, and Ec19 comparative genomics analysis**

**Figure S5 Comparison network diagram of the microcosm RNA-seq and TraDIS assays**

**Figure S6 Freshwater phage microbiome and validation and phenotypic characterisation of the Ec19 mutants**

**Supplemental Note**

### Supplementary Figures

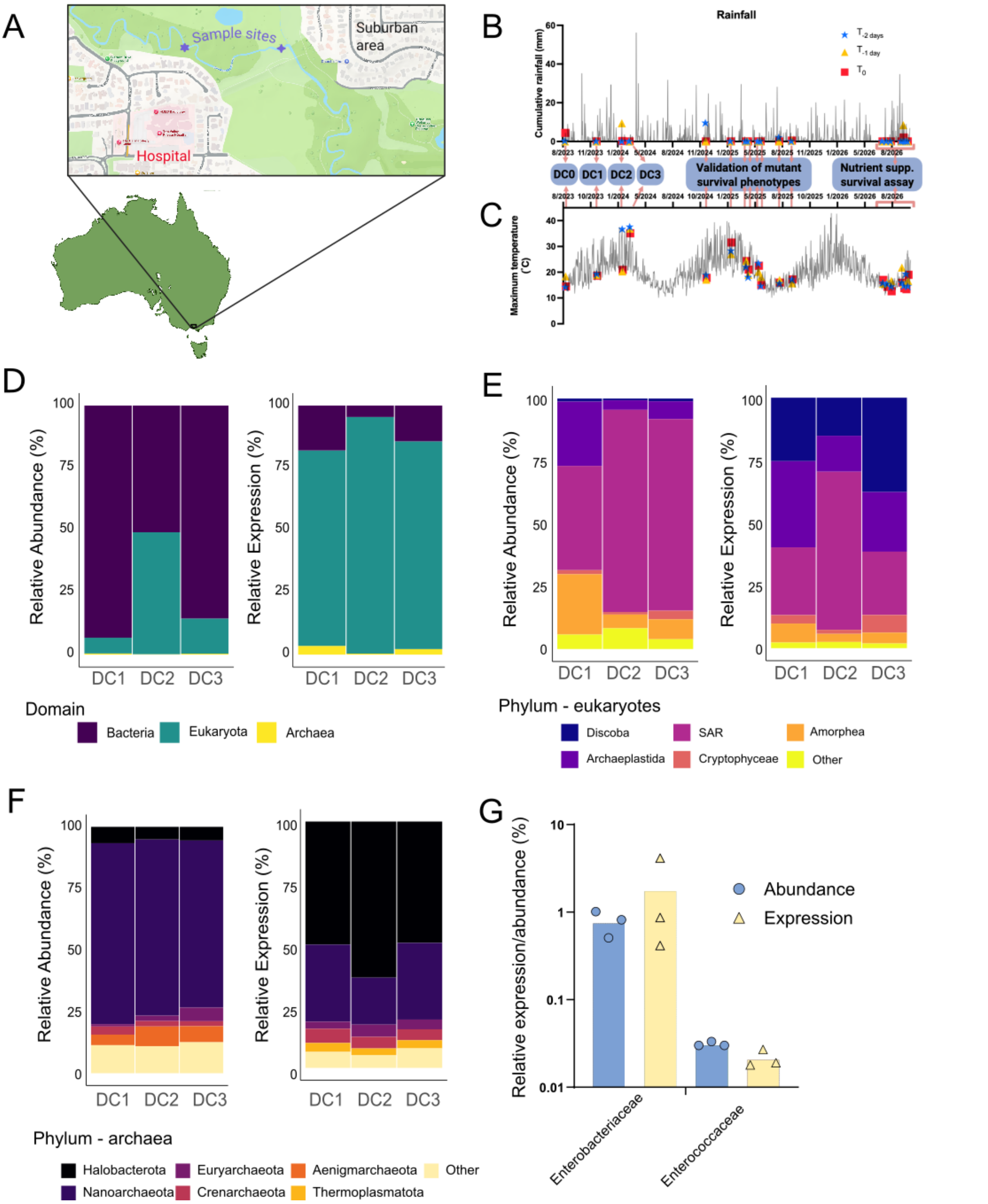


**Figure S1 Sampling sites and freshwater microbial community and target genera abundance and expression.** A) Sampling sites used for Enterobacteriaceae isolation and creek water collection for microcosm assays. The sampling site at coordinates -37.936845,

145.218046 (four-pointed star) was used to collect creek water for Metatranscriptomics, Microcosm RNA-seq and Microcosm TraDIS experiments, as well as for isolation of environmental Enterobacteriaceae. The sampling site at coordinates -37.936321, 145.214379 (six-pointed star) was used to collect creek water for Microcosm survival assays. B) Daily cumulative rainfall of the Dandenong Creek sampling site between May 2023 and September 2025. Rainfall data was obtained from a publicly available rainfall database in Melbourne Water (https://www.melbournewater.com.au/water-and-environment/water-management/rainfall-and-river-levels#/). C) Daily maximum temperature of Melbourne between May 2023 and September 2025. Temperature data was obtained from a publicly available database in the Bureau of Meteorology (https://www.bom.gov.au/climate/data/stations/). The corresponding rainfall and temperature two days before ($T_{-2\ days}$), one day before ($T_{-1\ day}$) and the day of the creek water collection ($T_0$) are highlighted in B) and C). Relative abundance and expression of each domain (D), top eukaryote (E) and archaea phyla (F) determined by phyloflash, and relative abundance of target genera (G) determined via kraken2.

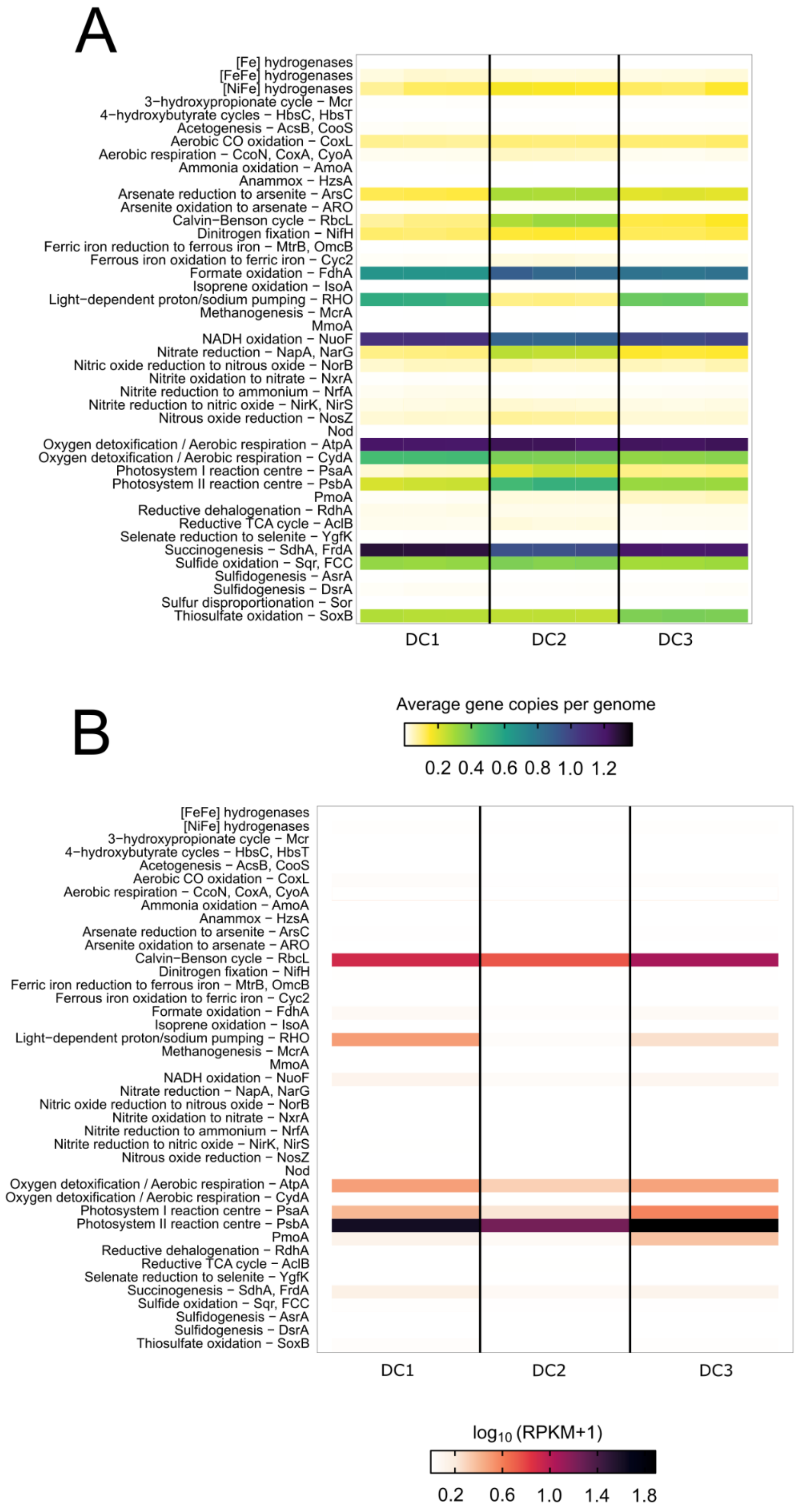

A
[Fe] hydrogenases
[FeFe] hydrogenases
[NiFe] hydrogenases
3−hydroxypropionate cycle − Mcr
4−hydroxybutyrate cycles − HbsC, HbsT
Acetogenesis − AcsB, CooS
Aerobic CO oxidation − CoxL
Aerobic respiration − CcoN, CoxA, CyoA
Ammonia oxidation − AmoA
Anammox − HzsA
Arsenate reduction to arsenite − ArsC
Arsenite oxidation to arsenate − ARO
Calvin−Benson cycle − RbcL
Dinitrogen fixation − NifH
Ferric iron reduction to ferrous iron − MtrB, OmcB
Ferrous iron oxidation to ferric iron − Cyc2
Formate oxidation − FdhA
Isoprene oxidation − IsoA
Light−dependent proton/sodium pumping − RHO
Methanogenesis − McrA
MmoA
NADH oxidation − NuoF
Nitrate reduction − NapA, NarG
Nitric oxide reduction to nitrous oxide − NorB
Nitrite oxidation to nitrate − NxrA
Nitrite reduction to ammonium − NrfA
Nitrite reduction to nitric oxide − NirK, NirS
Nitrous oxide reduction − NosZ
Nod
Oxygen detoxification / Aerobic respiration − AtpA
Oxygen detoxification / Aerobic respiration − CydA
Photosystem I reaction centre − PsaA
Photosystem II reaction centre − PsbA
PmoA
Reductive dehalogenation − RdhA
Reductive TCA cycle − AclB
Selenate reduction to selenite − YgfK
Succinogenesis − SdhA, FrdA
Sulfide oxidation − Sqr, FCC
Sulfidogenesis − AsrA
Sulfidogenesis − DsrA
Sulfur disproportionation − Sor
Thiosulfate oxidation − SoxB
DC1
DC2
DC3
Average gene copies per genome
0.2 0.4 0.6 0.8 1.0 1.2
B
[FeFe] hydrogenases
[NiFe] hydrogenases
3−hydroxypropionate cycle − Mcr
4−hydroxybutyrate cycles − HbsC, HbsT
Acetogenesis − AcsB, CooS
Aerobic CO oxidation − CoxL
Aerobic respiration − CcoN, CoxA, CyoA
Ammonia oxidation − AmoA
Anammox − HzsA
Arsenate reduction to arsenite − ArsC
Arsenite oxidation to arsenate − ARO
Calvin−Benson cycle − RbcL
Dinitrogen fixation − NifH
Ferric iron reduction to ferrous iron − MtrB, OmcB
Ferrous iron oxidation to ferric iron − Cyc2
Formate oxidation − FdhA
Isoprene oxidation − IsoA
Light−dependent proton/sodium pumping − RHO
Methanogenesis − McrA
MmoA
NADH oxidation − NuoF
Nitrate reduction − NapA, NarG
Nitric oxide reduction to nitrous oxide − NorB
Nitrite oxidation to nitrate − NxrA
Nitrite reduction to ammonium − NrfA
Nitrite reduction to nitric oxide − NirK, NirS
Nitrous oxide reduction − NosZ
Nod
Oxygen detoxification / Aerobic respiration − AtpA
Oxygen detoxification / Aerobic respiration − CydA
Photosystem I reaction centre − PsaA
Photosystem II reaction centre − PsbA
PmoA
Reductive dehalogenation − RdhA
Reductive TCA cycle − AclB
Selenate reduction to selenite − YgfK
Succinogenesis − SdhA, FrdA
Sulfide oxidation − Sqr, FCC
Sulfidogenesis − AsrA
Sulfidogenesis − DsrA
Thiosulfate oxidation − SoxB
DC1
DC2
DC3
log10 (RPKM+1)
0.2 0.6 1.0 1.4 1.8

**Figure S2. Metabolic, antibiotic resistance, virulence gene abundance and expression in freshwater microbial communities and isolates.** A) diamond screen of metabolic marker genes in metagenome reads from triplicate urban freshwater samples - abundance normalised to ribosomal marker genes and B) diamond screen of metabolic marker genes in metatranscriptome samples.

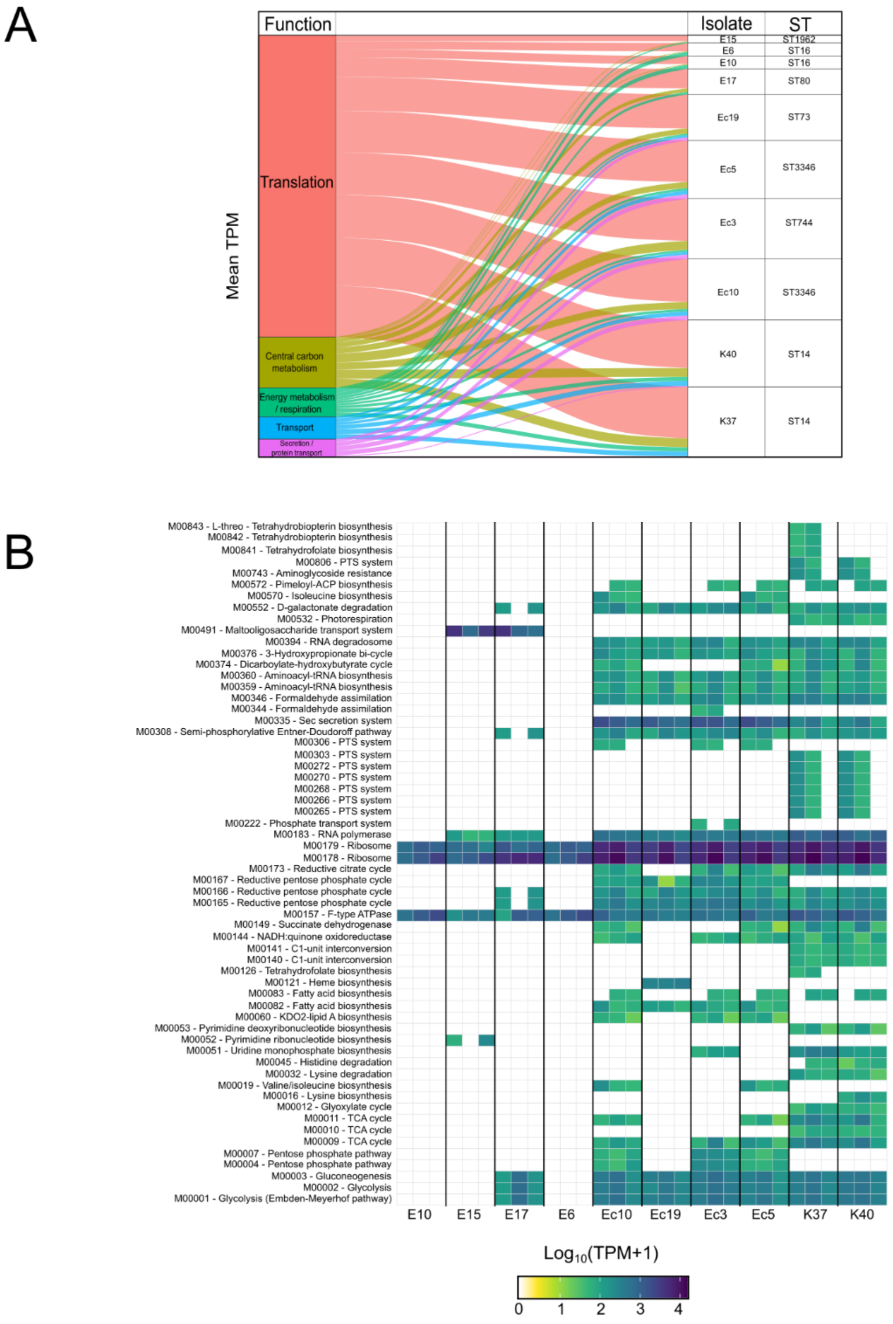


**Figure S3. COG categories and KEGG modules actively expressed by isolates *in situ.*** A) Most prominent *in situ* expression of genes specific to each isolate collapsed to COG (Cluster of Orthologous Genes) averaged from three independent surface water samples. B) *In situ* expression of genes in each isolate grouped by KEGG modules, the data was filtered to focus the analysis using a threshold of >50 TPM, and present in at least two replicates.

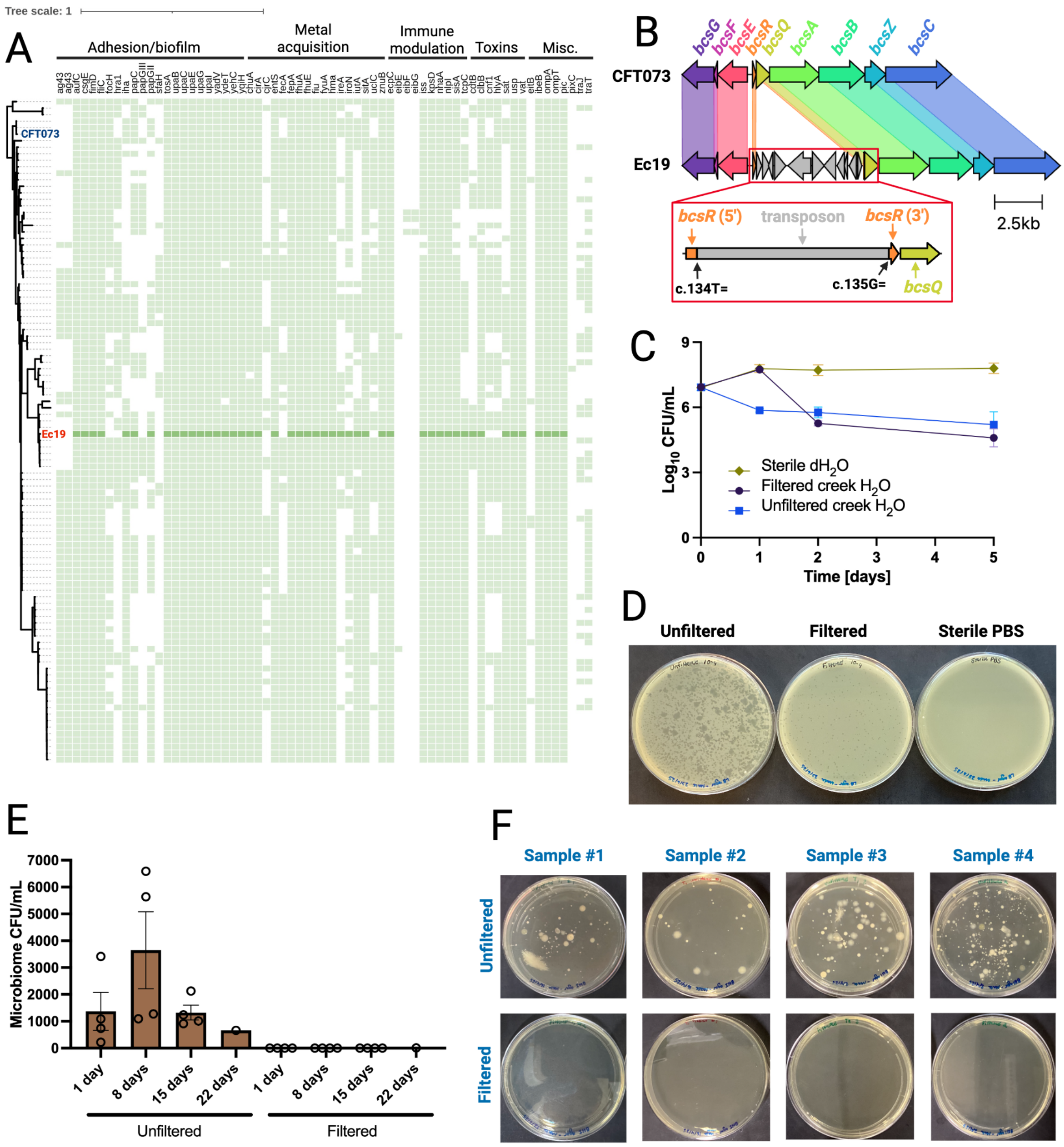


**Figure S4 Creek water survival, microbiome, and Ec19 comparative genomics analysis.** A) Phylogenetic tree and presence-absence matrix of virulence factors of *E. coli* Ec19, CFT073 and 100 randomly selected ST73 isolates from Enterobase. A) Phylogenetic tree comparing Ec19 and 50 pathogenic and commensal E. coli strains. The corresponding sequence type and pathotype of each strain are shown in blue. The presence-absence matrix of virulence and antibiotic resistance genes for each strain are shown in green and pink respectively, with the Ec19-associated genes shown as darker variants of those colours. BB) Comparison of the cellulose (bcs) operon between a cellulose-positive (CFT073) and cellulose-negative (Ec19) E. coli. Ec19 possesses a transposon inserted between the 134th and 135th nucleotide position of *bcsR*. C) Ec19 survival curves in various water conditions over a 5-day period. Each data point represents the average CFU across three biological and two technical replicates, while the error bars represent the standard error of the mean. D) LB top agar inoculated with unfiltered or filtered water or sterile PBS. E) The average microbiome content, expressed in CFU/mL, of filtered and unfiltered creek water samples after one, eight, 15 and 22 days of storage at 4˚C. FE) Brain Heart Infusion agar plates spread plated with filtered or unfiltered 1-day-old creek water samples collected across different sampling dates.

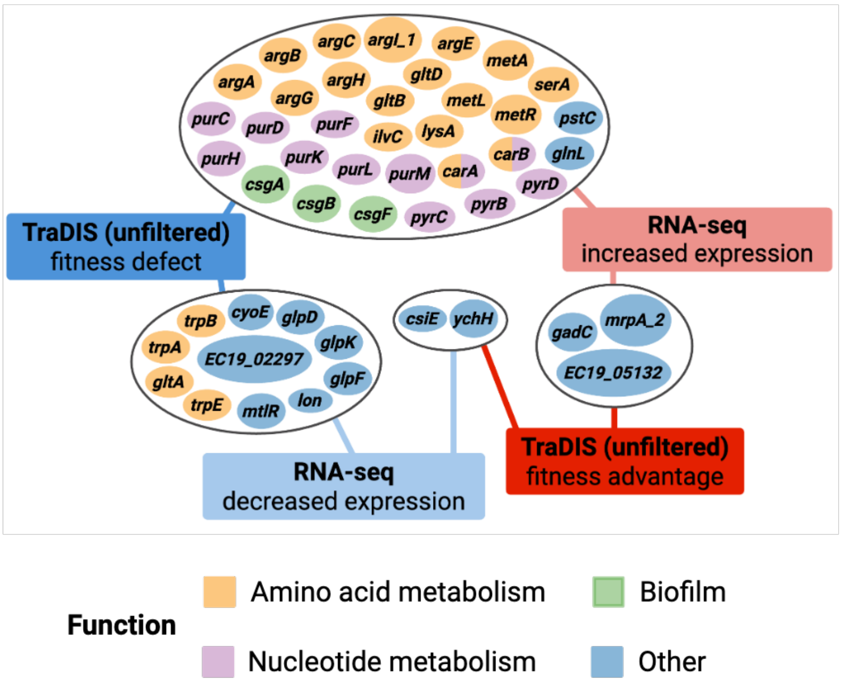


**Figure S5 Comparison network diagram of the microcosm RNA-seq and TraDIS assays.** This diagram illustrates all Ec19 genes that were significant hits in both RNA-seq and TraDIS experiments.

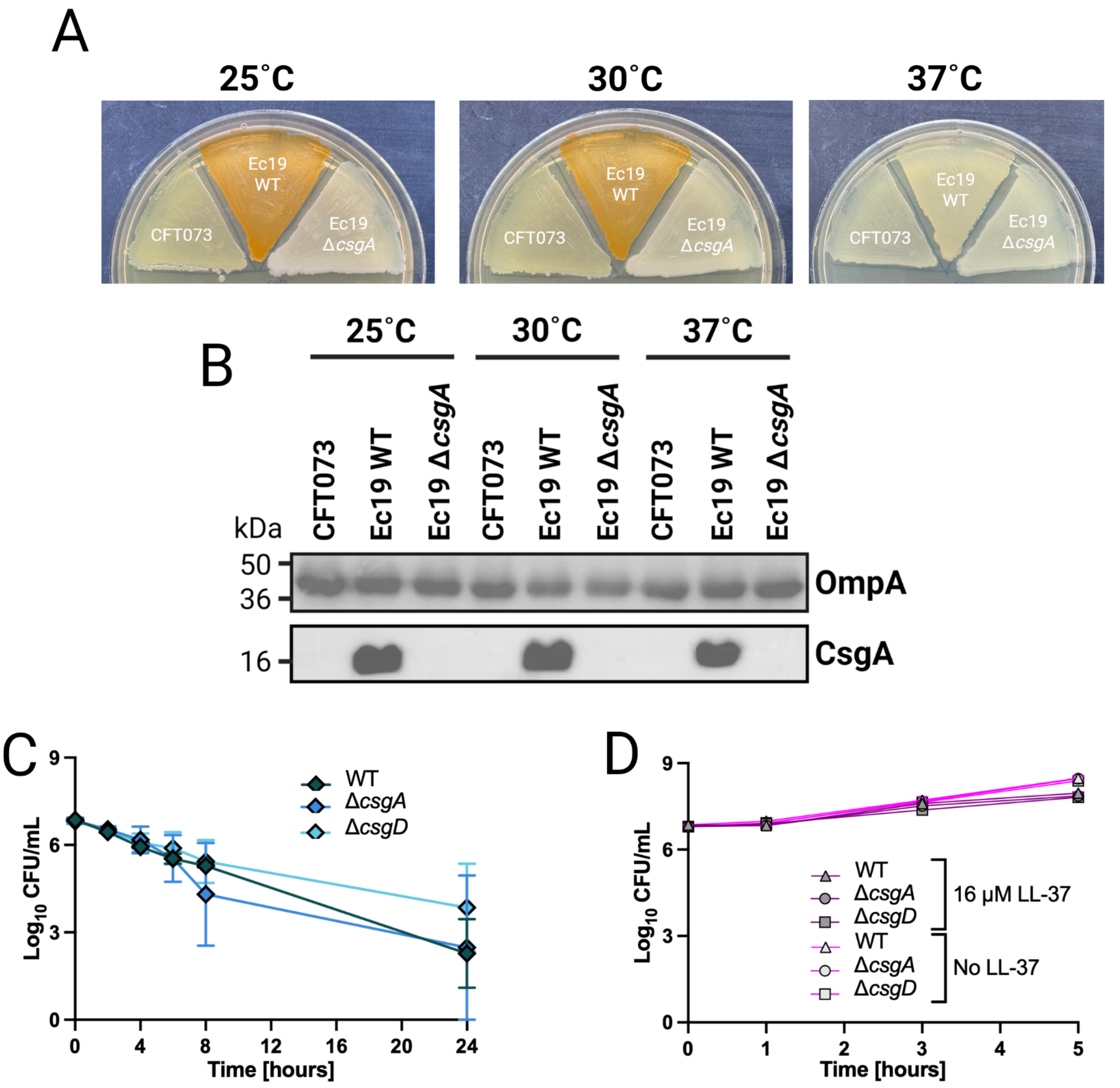


**Figure S6 Creek water phage microbiome and validation and phenotypic characterisation of the Ec19 mutants.** A) Curcumin agar assay of *E. coli* Ec19 WT, Ec19Δ*csgA*, Ec19Δ*csgD*, CFT073 and BW25113 grown in 25˚C, 30˚C and 37˚C. B) Western blotting of CsgA and OmpA (loading control) from whole cell lysates of CFT073, Ec19 WT, Ec19Δ*csgA* and Ec19Δ*csgD* grown at 25˚C, 30˚C and 37˚C. C) Time-killing curve of Ec19 WT and curli mutants exposed to 3 µg/mL Polymyxin B over a 24-hour period. D) Time-killing curve of Ec19 WT and curli mutants with or without exposure to 16 µM LL-37 over a 5-hour period. Each data point in CB) and DC) represent the average value across two biological and two technical replicates, and the error bars represent the standard error of the mean.

**Supplemental Note**

Isolate pathways actively expressed *in situ*

All isolates showed active expression of RNA polymerase (M00183), ribosome (M00178, M00179), F-type ATPase (M00157) and variably glycolysis and gluconeogenesis (M00001, M00002, M00003). *Enterococcus* spp. isolates showed very few active processes, where *E. faecium* strain E15 showed active expression of pyrimidine ribonucleotide biosynthesis (M00052) and both *E. faecium* isolates E15 and E17 showed strong expression of malto-oligosaccharide transport systems (M00491) (Figure 2B, Figure S3B). By contrast, Enterobacteriaceae isolates showed a multitude of active metabolic processes including carbon metabolism (glycolysis, gluconeogenesis, pentose phosphate pathway, tri-carboxylic acid (TCA) cycle, D-galactonate degradation, pyruvate oxidation), respiration (NADH:quinone oxidoreductase/complex I, succinate dehydrogenase/complex II, F-type ATPase, cytochrome bd oxidase), amino acid biosynthesis, degradation and transport, phosphoenolpyruvate-dependent sugar phosphotransferase (PTS) systems, Sec secretion, RNA degradosome, and aminoglycoside resistance (Figure 2B, Figure S2B). The widespread expression of D-galactonate degradation suggests this metabolite could represent a significant carbon source within freshwater, and genes dedicated to D-galactonate catabolism have previously been identified in the genomes of freshwater bacteria and environmental Salmonella enterica isolates (1). Additionally, only few carbohydrate active enzymes (CAZymes) were expressed in Enterobacteriaceae *in situ*, including genes encoding putative amidases (*bax*), endoglucanases (*bcsZ*), lipid A modification enzymes (*arnC*) and maltodextrin glucosidases (*malZ*). Enterobacteriaceae isolates also showed widespread expression of amino acid metabolism (degradation, transport and biosynthesis). Finally, the siderophore Aerobactin (*iucC*) was expressed in all *E. coli* similar to the whole community analysis, these data together suggest that suggesting that these bacteria are utilising multiple strategies to meet their energy and nutrient requirements in the nutrient limited freshwater environment.

1. Aguirre-Sanchez JR, Ibarra-Rodriguez JR, Vega-Lopez IF, Martínez-Urtaza J, Chaidez-Quiroz C. 2021. Genomic signatures of adaptation to natural settings in non-typhoidal Salmonella enterica Serovars Saintpaul, Thompson and Weltevreden. Infect Genet Evol 90:104771.
